# Genome sequencing and analysis of the first spontaneous Nanosilver resistant bacterium *Proteus mirabilis* strain SCDR1

**DOI:** 10.1101/089961

**Authors:** Amr T. M. Saeb, Khalid A. Al-Rubeaan, Mohamed Abouelhoda, Manojkumar Selvaraju, Hamsa T. Tayeb

## Abstract

**Background:** *P. mirabilis* is a common uropathogenic bacterium that can cause major complications in patients with long-standing indwelling catheters or patients with urinary tract anomalies. In addition, *P. mirabilis* is a common cause of chronic osteomyelitis in Diabetic foot ulcer (DFU) patients. We isolated *P. mirabilis SCDR1* from a Diabetic ulcer patient. We examined *P. mirabilis* SCDR1 levels of resistance against Nano-silver colloids, the commercial Nano-silver and silver containing bandages and commonly used antibiotics. We utilized next generation sequencing techniques (NGS), bioinformatics, phylogenetic analysis and pathogenomics in the characterization of the infectious pathogen.

**Results:** *P. mirabilis SCDR1* is a multi-drug resistant isolate that also showed high levels of resistance against Nano-silver colloids, Nano-silver chitosan composite and the commercially available Nano-silver and silver bandages. The *P. mirabilis* -SCDR1 genome size is 3,815,621 bp. with G+C content of 38.44%. *P. mirabilis*-SCDR1 genome contains a total of 3,533 genes, 3,414 coding DNA sequence genes, 11, 10, 18 rRNAs (5S, 16S, and 23S), and 76 tRNAs. Our isolate contains all the required pathogenicity and virulence factors to establish a successful infection. *P. mirabilis* SCDR1 isolate is a potential virulent pathogen that despite its original isolation site, wound, it can establish kidney infection and its associated complications. *P. mirabilis SCDR1* contains several mechanisms for antibiotics and metals resistance including, biofilm formation, swarming mobility, efflux systems, and enzymatic detoxification.

**Conclusion:** *P. mirabilis SCDR1* is the first reported spontaneous Nanosilver resistant bacterial strain. *P. mirabilis* SCDR1 possesses several mechanisms that may lead to the observed Nanosilver resistance.

## Background

The production and utilization of nanosilver are one of the primary and still growing application in the field of nanotechnology. Nanosilver is used as the essential antimicrobial ingredient in both clinical and environmental technologies. (Chen and Schluesener 2008; Franci et al. 2015; Oyanedel-Craver and Smith 2008; Prabhu and Poulose 2012). Nanosilver is known to exert inhibitory and bactericidal effects activities against many Gram-positive, Gram-negative and fungal pathogens (Saeb et al. 2014). Latest studies suggest that the use of nanosilver-containing wound dressings prevent or reduce microbial growth in wounds and may improve the healing process (Velázquez-Velázquez et al. 2015). Moreover, antibacterial nanosilver-containing wound dressing gels may be important for patients that are at risk for non-healing of diabetic foot wounds and traumatic/surgical wounds (Lullove and Bernstein 2015). Increased usage of nanosilver in both medical and environmental products has generated concerns about the development of bacterial resistance against the antimicrobial ingredient. Bacterial resistance against metallic silver has been documented several bacterial strains such as *E. coli Enterobacter cloacae, Klebsiella pneumoniae* and *Salmonella typhimurium* (Hendry and Stewart 1979; McHugh et al. 1975). However, information about bacterial resistance against Nanosilver is in scarce. Only Gunawan et al., (2013) reported the occurrence of induced adaptation, of non-targeted environmental *Bacillus* species, to antimicrobial Nanosilver (Gunawan et al. 2013). In this study we are presenting *of Proteus mirabilis SCDR1* isolate, the first reported spontaneous Nanosilver resistant bacterial strain.

*Proteus mirabilis* is a motile gram-negative bacterium that is characterized by it swarming behavior (Jansen et al. 2003). Although it resides in human gut commensally, *P. mirabilis* is a common uropathogen that can cause major complications. In addition, *P. mirabilis* can cause respiratory and wound infections, bacteremia, and other infections (Mathur et al. 2005; Armbruster and Mobley 2012; Jacobsen et al. 2008). In fact, *P. mirabilis* is a common cause of chronic osteomyelitis in Diabetic foot ulcer (DFU) patients (Bronze and Cunha 2016). Generally, *P. mirabilis* is responsible for 90% of genus *Proteus* infections and can be considered as a community-acquired infection (Gonzalez and Bronze 2016). As a pathogen *P. mirabilis* acquires many virulence determinants that enable it to establish successful infections. Alongside with mobility (flagellae), adherence, hemolysin, toxin production, Urease, Quorum sensing, iron acquisition systems, and proteins that function in immune evasion, are important virulence factors of *P. mirabilis* (Habibi et al. 2015; Baldo and Rocha 2014). A lot of information concerning antibiotic resistance are available for *P. mirabilis* (Horner et al. 2014; Miró et al. 2013; Hawser et al. 2014). *P. mirabilis* is intrinsically resistant to tetracyclines and polymyxins. Moreover, multidrug-resistant (MDR) *P. mirabilis* strains resistant resistance to β-lactams, aminoglycosides, fluoroquinolones, phenicols, streptothricin, tetracycline, and trimethoprim-sulfamethoxazole was reported (Chen et al. 2015). However, limited information about heavy metals, including silver, is available. In this study, we are presenting the first report and genome sequence of nanosilver resistant bacterium *P. mirabilis* strain SCDR1 isolated from diabetic foot ulcer (DFU) patient.

## Materials and Methods

### Bacterial isolate

*Proteus mirabilis* strain SCDR1 was isolated from a Diabetic ulcer patient in the Diabetic foot unit in the University Diabetes Center at King Saud University. Proper wound swab was obtained from the patient and was sent for further microbiological study and culture. Wounds needing debridement were debrided before swabbing the surface of the wound. Specimen was inoculated onto blood agar (BA; Oxoid, Basingstoke, UK) and MacConkey agar (Oxoid) and incubated at 37°C for 24 - 48 h. The Vitek 2 system and its advanced expert system were used for microbial identification, antibiotic sensitivity testing, and interpretation of results. Manual disk diffusion and MIC method for AgNPs and antibiotic sensitivity testing were performed when required.

### Preparation of colloidal and composite Nanosilver and Commercial products for antimicrobial activity testing

Colloidal silver Nanoparticles were prepared, characterized and concentration determined as described by Saeb et al., 2014 (Saeb et al. 2014). Nanosilver chitosan composite preparations were done by chemical reduction method as described by Latif et al., 2015 (Latif et al. 2015). Moreover, the following commercially silver and Nanosilver containing wound dressing bandages were also used for antimicrobial activity testing: Silvercel non-adherent antimicrobial alginate Dressing (Acelity L.P. Inc, San Antonio, Texas, USA), Sorbsan Silver dressing made of Calcium alginate with silver (Aspen Medical Europe Ltd., Leicestershire, UK), ColActive^®^ Plus Ag (Covalon Technologies Ltd., Mississauga, Ontario, Canada), exsalt^®^SD7 wound dressing (Exciton Technologies, Edmonton, Alberta, Canada), Puracol Plus AG+ Collagen Dressings with Silver (Medline, Mundelein, Illinois, USA) and ACTISORB^™^ silver antimicrobial wound dressing 220 (Acelity L.P. Inc, San Antonio, Texas, USA).

### Antimicrobial Susceptibility Test

Antimicrobial activities were performed against the following strains: *Pseudomonas aeruginosa* ATCC 27853, *Staphylococcus aureus* ATCC 29213, *Proteus mirabilis* ATCC 29906, *Klebsiella pneumoniae* ATCC 700603, *E. coli* ATCC 25922 and *Enterobacter cloacae* ATCC 29212.

### Disk diffusion antimicrobial susceptibility testing

Disk diffusion antimicrobial susceptibility testing was performed as described by Matuschek et al. (Matuschek et al. 2014). The sterile discs were loaded with different concentrations (50-200 ppm) of colloidal silver nanoparticles solutions and the Nanosilver chitosan composite (composite concentration ranged between 0.1% and 0.01M to 3.2% and 0.16M from chitosan and Silver nitrate respectively) and then placed on Mueller–Hinton (MH) agar plates with bacterial lawns. Within 15 min of application of antimicrobial disks, the plates were inverted and incubated 37°C for 16 hours. All experiments were done in an aseptic condition in laminar air flow cabinet. After incubation, inhibition zones were read at the point where no apparent growth is detected. The inhibition zone diameters were measured to the nearest millimeter. Similarly, 5mm desks from the commercially available bandages were prepared in an aseptic condition and tested for their antimicrobial activity as described before.

### Minimum bactericidal (MBC), Minimal inhibitory concertation (MIC) and Biofilm formation tests

MBC and MIC testing were performed as described by Holla et al., (Holla et al. 2012). Different volumes that contained a range of silver Nanoparticles (50-700 ppm) were delivered to 7.5 ml of Muller-Hinton (MH) broth each inoculated with 0.2 ml of the bacterial suspensions. Within 15 min of application of silver nanoparticles, the tubes were incubated at 37°C for 16 hours in a shaker incubator at 200 rpm. We included a positive control (tubes containing inoculum and nutrient media without silver nanoparticles) and a negative control (tubes containing silver nanoparticles and nutrient media without inoculum).Biofilm formation ability of *P. mirabilis* SCDR1 was tested as described before by Yassien and Khardori (Yassien and Khardori 2001).

## Molecular Genomics analysis

### DNA purification and Sequencing

Maxwell 16 automated DNA isolation machine was used for DNA isolation according to the instructions of the manufacturer. Isolated DNA was quantified using NanoDrop 2000c UV-Vis spectrophotometer. The Agilent 2100 Bioanalyzer system will be used for sizing, quantitation and quality control of DNA. The quality of subjected DNA sample was determined by loading a 150 mg of diluted DNA in 1% agarose E-gel (Invitrogen, Paisley, UK). We have conducted two sequencing runs using the Personal Genome Machine (PGM) sequencer from Life Technologies (Thermo Fischer) according to the instructions of the manufacture.

### Bioinformatics analysis

We have developed an analysis pipeline to identify the suggested pathogen and annotate it. First, the quality of the reads was assessed and reads with a quality score less than 20bp were trimmed out. The reads were then passed to the program Metaphlan (Segata et al. 2012) for primary identifications of microbial families included in the samples based on unique and clade-specific marker genes. In parallel to run Metaphlan analysis, we used BLAST program to map each read to the non-redundant nucleotide database of NCBI. We mapped the reads back to the bacterial genomes thought to be the pathogen; these are the top ranked bacteria based on Metaphlan, BLAST results, and related taxa analysis. The integration of the different tools and execution of the whole pipeline is achieved through python scripts developed in-house. A version of this pipeline is currently being imported to the workflow system Tavaxy (Abouelhoda et al. 2012) to be used by other researchers. Furthermore, we retrieved the genome annotation from the Genbank and investigated the missing genes. In addition, we used QIIME the open-source bioinformatics pipeline for performing microbiome analysis from raw DNA sequencing data for taxonomic assignment and results visualizations (Caporaso et al. 2010).

### Phylogenetic analysis

The 16S rDNA sequences of our isolate were used to construct a phylogenetic relationship with other *Proteus mirabilis* species. We acquired partial 16S rDNA sequences of selected *Proteus mirabilis* species from the GenBank. In order to establish the phylogenetic relationships among taxa, phylogenetic trees were constructed using the Maximum Likelihood (ML) method based on the Jukes-Cantor model the best fit to the data according to AIC criterion (Tamura and Nei 1993). MEGA6 (program / software/tool) was used to conduct phylogenetic analysis (Tamura et al. 2013, 0). In addition, a whole genome Neighbor-joining phylogenetic distance based tree *of Proteus mirabilis* spices including *Proteus mirabilis* SCDR1 isolate using the BLAST new enhanced graphical presentation and added functionality available at https://blast.ncbi.nlm.nih.gov/ (National Center for Biotechnology Information). In addition, we used Mauve (Darling et al. 2004) and CoCoNUT (Abouelhoda et al. 2008) to generate the whole genome pairwise and multiple alignments of the draft *P. mirabilis* strain SCDR1 genome against selected reference genomes. Furthermore, we performed whole genome phylogeny based proteomic comparison among *P. mirabilis* SCDR1 isolate and other related *Proteus mirabilis* strains using Proteome Comparison service which is protein sequence-based comparison using bi-directional BLASTP available at (https://www.patricbrc.org/app/SeqComparison) (Wattam et al. 2014).

### Gene annotation and Pathogenomics analysis

*P. mirabilis* SCDR1 genome contigs were annotated using the Prokaryotic Genomes Automatic Annotation Pipeline (**PGAAP**) available at NCBI (http://www.ncbi.nlm.nih.gov/). In addition, contigs were further annotated using the bacterial bioinformatics database and analysis resource (PATRIC) gene annotation service (https://www.patricbrc.org/app/Annotation) (Wattam et al. 2014). The **PathogenFinder 1.1** pathogenicity prediction program available at (https://cge.cbs.dtu.dk/services/PathogenFinder/) was used to examine the nature of *P. mirabilis* SCDR1 as a human pathogen (Cosentino et al. 2013). Virulence genes sequences and functions, corresponding to different major bacterial virulence factors *of Proteus mirabilis* were collected from GenBank and validated using virulence factors of pathogenic bacteria database available at (http://www.mgc.ac.cn/VFs/) (2003), Victors virulence factors search program available at (http://www.phidias.us/victors/) and PATRIC_VF tool available at https://www.patricbrc.org/portal/portal/patric/SpecialtyGeneSource?source=PATRIC_VF&kw= (Wattam et al. 2014).

### Resistome analysis

*P. mirabilis* SCDR1 genome contigs were investigated manually for the presence of antibiotic resistance loci using **PGAAP** and **PATRIC** gene annotation services. Antibiotic resistance loci were further investigated using specialized search tools and services namely, **Antibiotic Resistance Gene Search** available at (https://www.patricbrc.org/portal/portal/patric/AntibioticResistanceGeneSearch?cType=taxon&cId=131567&dm=) (Wattam et al. 2014), **Genome Feature Finder** (antibiotic resistance) available at (https://www.patricbrc.org/portal/portal/patric/GenomicFeature?cType=taxon&cId=131567&dm=) (Wattam et al. 2014), **ARDB** (Antibiotic Resistance Genes Database) available at https://ardb.cbcb.umd.edu/) (Liu and Pop 2009), **CARD** (The Comprehensive Antibiotic Resistance Database) available at (https://card.mcmaster.ca/) (McArthur and Wright 2015; McArthur et al. 2013), **Specialty Gene Search** available at https://www.patricbrc.org/portal/portal/patric/SpecialtyGeneSearch?cType=taxon&cId=131567&dm=) and **ResFinder 2.1** available at https://cge.cbs.dtu.dk//services/ResFinder/ (Zankari et al. 2012).

The heavy metal resistance gene search *P. mirabilis* SCDR1 contigs were investigated using **PGAAP** and **PATRIC** gene annotation services, **PATRIC Feature Finder** searches tool and **BacMet** (antibacterial biocide and metal resistance genes database) available at (http://bacmet.biomedicine.gu.se/) (Wattam et al. 2014; Pal et al. 2014).

## Results

### Initial identification and Antimicrobial Susceptibility Test

The Vitek 2 system showed that our isolate belongs to *Proteus mirabilis* species. Antibiotic sensitivity testing using Vitek 2 AST-N204 card showed that our isolate *P. mirabilis* SCDR1 is resistant to ampicillin, nitrofurantoin, and Trimethoprim/ Sulfamethoxazole. In addition, *P. mirabilis* SCDR1 was resistant against ethidium Bromide, tetracycline, tigecycline, colistin, polymyxin B, rifamycin, doxycycline, vancomycin, fusidic acid, bacitracin, metronidazole, clarithromycin, erythromycin, oxacillin, clindamycin, trimethoprim, novobiocin, and minocycline. *P. mirabilis* SCDR1 was intermediate resistant against nalidixic acid, Imipenem, and Cefuroxime. Whereas it was sensitive to chloramphenicol, amoxicillin/ clavulanic Acid, piperacillin/tazobactam, cefotaxime, ceftazidime, cefepime, cefaclor, cephalothin, ertapenem, meropenem, amikacin, gentamicin, ciprofloxacin, norfloxacin, tobramycin, streptomycin, and fosfomycin.

*P. mirabilis* SCDR1 isolate showed high resistance against colloidal and composite Nanosilver and metallic silver compared with other tested Gram positive and negative bacterial species. For instance, Table 1, shows the resistance of **P. mirabilis* SCDR1* against colloidal Nanosilver assessed by disk diffusion method in comparison with *S. aureus* ATCC 29213, *P. aeruginosa* ATCC 27853, *E. coli* ATCC 25922 and *E. cloacae* ATCC 29212. Generally, **P. mirabilis* SCDR1* showed high resistance (0.0 cm), while *K. pneumoniae* showed the highest sensitivity (1.5-1.9 cm) against all tested silver nanoparticle concentrations (50-200 ppm). *S. aureus* also showed high sensitivity (1.4-1.6 cm) against all tested silver nanoparticle concentrations. None of the tested bacterial isolates, except for **P. mirabilis* SCDR1,* showed any resistance against silver-nanoparticles even against the lowest concentration (50 ppm).

**Table 1:** Resistance of *P. mirabilis* SCDR1 against colloidal Nano-Silver assessed by desk diffusion method.

| S. No. | Sample ID | Zone Of Inhibition (cm)<br>*S. aureus* | Zone Of Inhibition (cm)<br>*E. cloacae* | Zone Of Inhibition (cm)<br>*P. aeruginosa* | Zone Of Inhibition (cm)<br>*E. coli* | Zone Of Inhibition (cm)<br>*K. pneumoniae* | Zone Of Inhibition (cm)<br>*P. mirabilis* SCDR1 |
| --- | --- | --- | --- | --- | --- | --- | --- |
| 1 | 200 ppm | 1.6 cm | 1.5 cm | 1.4 cm | 1.1 cm | 1.9 cm | 0.0 cm |
| 2 | 150 ppm | 1.5 cm | 1.2 cm | 1.3 cm | 1.0 cm | 1.7 cm | 0.0 cm |
| 3 | 100 ppm | 1.5 cm | 1.2 cm | 1.3 cm | 1.0 cm | 1.6 cm | 0.0 cm |
| 4 | 50 ppm | 1.4 cm | 1.1 cm | 1.1 cm | 0.9 cm | 1.5 cm | 0.0 cm |

Furthermore, Table 2 shows the resistance of *P. mirabilis* SCDR1 against colloidal Nanosilver assessed by minimal inhibitory concentration method compared with other tested Gram positive and negative bacterial species. Once more, **P. mirabilis* SCDR1* showed high resistance against the gradually increased concentrations of colloidal Nano-Silver. We observed **P. mirabilis* SCDR1* bacterial growth to colloidal Nanosilver concentration up to 500 ppm. On the other hand, *K. pneumoniae* showed the highest sensitivity against silver nanoparticles with no observed growth at only 100 ppm colloidal Nanosilver concentration. In addition, both *E. coli* and *P. aeruginosa* showed the high sensitivity against silver nanoparticles with no observed growth at 150 ppm colloidal Nanosilver concentration. While, *S.aureus* tolerated only 200 ppm colloidal Nanosilver concentration.

**Table 2:** Resistance of *P. mirabilis* SCDR1 against colloidal Nanosilver assessed by minimal inhibitory concentration method.

| AgNPs<br>(concentration in<br>ppm) | Bacterial species/strain |  |  |  |  |  |  |
| --- | --- | --- | --- | --- | --- | --- | --- |
|  | *S. aureus*<br>ATCC 29213 | *P. aeruginosa*<br>ATCC 27853 | *E. cloacae*<br>ATCC 29212 | *E. coli*<br>ATCC<br>25922 | *K. pneumoniae*<br>ATCC 700603 | *P. mirabilis*<br>SCDR1 | *P. mirabilis*<br>ATCC 29906 |
| 50 | Growth | Growth | Growth | Growth | Growth | Growth | Growth |
| 100 | Growth | Growth | Growth | Growth | No Growth | Growth | Growth |
| 150 | Growth | No Growth | Growth | No Growth | No Growth | Growth | Growth |
| 200 | Growth | No Growth | Growth | No Growth | No Growth | Growth | Growth |
| 250 | No Growth | No Growth | No Growth | No Growth | No Growth | Growth | Growth |
| 300 | No Growth | No Growth | No Growth | No Growth | No Growth | Growth | Growth |
| 350 | No Growth | No Growth | No Growth | No Growth | No Growth | Growth | Growth |
| 400 | No Growth | No Growth | No Growth | No Growth | No Growth | Growth | Growth |
| 450 | No Growth | No Growth | No Growth | No Growth | No Growth | Growth | Growth |
| 500 | No Growth | No Growth | No Growth | No Growth | No Growth | Growth | No Growth |
| 550 | No Growth | No Growth | No Growth | No Growth | No Growth | No Growth | No Growth |
| 600 | No Growth | No Growth | No Growth | No Growth | No Growth | No Growth | No Growth |
| 650 | No Growth | No Growth | No Growth | No Growth | No Growth | No Growth | No Growth |
| 700 | No Growth | No Growth | No Growth | No Growth | No Growth | No Growth | No Growth |

Similarly, Table 3 shows the resistance of *P. mirabilis* SCDR1 against silver and Nanosilver composite assessed by disk diffusion method. Nanosilver chitosan composites, with concentration, ranged between 0.1% and 0.01M to 3.2% and 0.16M from chitosan and Silver nitrate respectively, had a comparable killing effect on both Gram positive and negative bacterial namely, *S. aureus* and *P. aeruginosa.* While none of the tested Nanosilver chitosan composites had any killing effect on *P. mirabilis* SCDR1. Similarly, all the tested commercially available silver and Nanosilver containing wound dressing bandages showed the enhanced killing effect on both *S. aureus* and *P. aeruginosa.* However, all these wound dressing bandages failed to inhibit *P. mirabilis* SCDR1 growth. *P. mirabilis* SCDR1 was able to produce strong biofilm with OD of 0.296.

**Table 3:** Resistance of *P. mirabilis* SCDR1 against silver and Nanosilver composite assessed by desk diffusion method.

| Sample ID | Zone Of Inhibition (cm) | Zone Of Inhibition (cm) | Zone Of Inhibition (cm) |
| --- | --- | --- | --- |
|  | *S. aureus* | *P. aeruginosa* | *P. mirabilis* SCDR1 |
| A | 0.9 cm | 0.8 cm | No. Inhibition |
| B | 0.9 cm | 0.9 cm | No. Inhibition |
| C | 0.8 cm | 0.9 cm | No. Inhibition |
| D | 0.8 cm | 0.9 cm | No. Inhibition |
| E | 0.9 cm | 0.9 cm | No. Inhibition |
| F | 0.8 cm | 0.8 cm | No. Inhibition |
| G | 0.7 cm | 0.7 cm | No. Inhibition |
| H | 0.9 cm | 0.9 cm | No. Inhibition |
| I | 0.9 cm | 1.0 cm | No. Inhibition |
| J | 0.9 cm | 1.0 cm | No. Inhibition |
| K | 0.8 cm | 0.6 cm | No. Inhibition |
| L | 0.8 cm | 0.8 cm | No. Inhibition |
| M | 0.9 cm | 0.8 cm | No. Inhibition |
| N | 0.9 cm | 0.9 cm | No. Inhibition |
| O | 1.0 cm | 0.9 cm | No. Inhibition |
| P | 0.8 cm | 0.8 cm | No. Inhibition |
| Q | 0.9 cm | 0.7 cm | No. Inhibition |
| R | 0.9 cm | 0.8 cm | No. Inhibition |
| S | 0.8 cm | 0.9 cm | No. Inhibition |
| T | 1.0 cm | 0.9 cm | No. Inhibition |
| U | 0.8 cm | 0.8 cm | No. Inhibition |
| V | 0.9 cm | 0.8 cm | No. Inhibition |
| W | 0.9 cm | 0.8 cm | No. Inhibition |
| X | 1.0 cm | 0.8 cm | No. Inhibition |
| Y | 0.8 cm | 0.8 cm | No. Inhibition |
| Z | 0.7 cm | 0.7 cm | No. Inhibition |
| A1 | 0.8 cm | 0.7 cm | No. Inhibition |
| B2 | 0.9 cm | 0.7 cm | No. Inhibition |
| C3 | 0.9 cm | 0.8 cm | No. Inhibition |
| D4 | 0.6 cm | NA | No. Inhibition |
| Silvercel | 1.3 cm | 1.4 cm | No. Inhibition |
| Sorbsan silver | 1.9 cm | 2.0 cm | No. Inhibition |
| Colactive® Plus Ag | 1.5cm | 2.0cm | No. Inhibition |
| Exsalt™ SD7 | 1.5cm | 1.5cm | No. Inhibition |
| Puracol® Plus Ag | 1.4cm | 2.0cm | No. Inhibition |
| Actisorb® Silver 220 | 0.9cm | 1.2cm | No. Inhibition |

### General genome features

Data from our draft genome of *P. mirabilis* SCDR1 was deposited in the NCBI-GenBank and was assigned accession number LUFT00000000. The *P. mirabilis* SCDR1 assembly resulted in 63 contigs, with an N50 contig size of 227,512 bp nucleotides, and a total length of 3,815,621 bp. The average G+C content was 38.44%. Contigs were annotated using the Prokaryotic Genomes Automatic Annotation Pipeline (PGAAP) available at NCBI (http://www.ncbi.nlm.nih.gov/) providing a total of 3,533 genes, 3,414 coding DNA sequence genes, 11, 10, 18 rRNAs (5S, 16S, and 23S), and 76 tRNAs. On the other hand, the bacterial bioinformatics database and analysis resource (PATRIC) gene annotation analysis showed that the number of the observed coding sequence (CDS) is 4423, rRNA is 10 and tRNA is 71. The unique gene count for the different observed metabolic pathways is 2585 (Figure 1).

**Figure 1:**
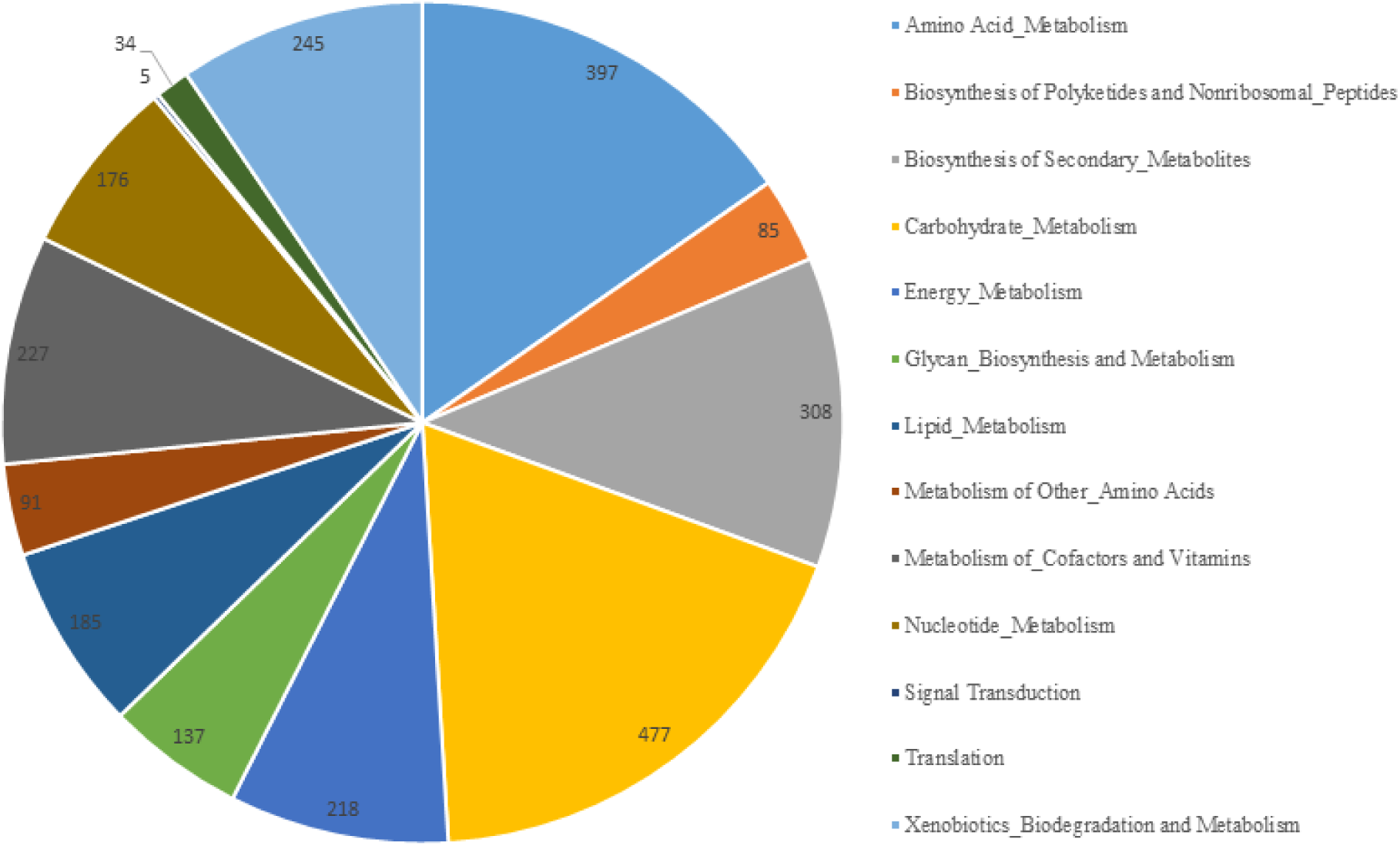
Distribution of unique gene counts amongst different metabolic pathways.

Carbohydrate metabolism pathways maintained the highest number of dedicated unique gene count (477) while signal transduction pathways maintained the highest number (5). In addition, biosynthesis of secondary metabolites, such as tetracycline, Streptomycin, Novobiocin, and Betalain, maintained a high number of dedicated unique gene (308). It is also noteworthy that Xenobiotics Biodegradation and Metabolism pathways also maintained a high number of dedicated unique gene (245) (**Supplementary table 1 and 2**).

### Pathogen identification and phylogenetic analysis

As stated before biochemical identification of the isolate confirmed the identity of our isolate to be belonging to *Proteus mirabilis* species. Moreover, Primary analysis of Metaphlan showed that *Proteus mirabilis* is the most dominant species in the sample (Figure 2). The appearance of other bacterial species in the Metaphlan diagram is explained by genomic homology similarity of other bacteria to *Proteus mirabilis. *P. mirabilis** SCDR1 genome showed high similarly 92.07% to the genome of *P. mirabilis* strain BB2000 followed by *P. mirabilis* strain C05028 (90.99%) and *P. mirabilis* strain PR03 (89.73%) (Table 4).

**Table 4:**
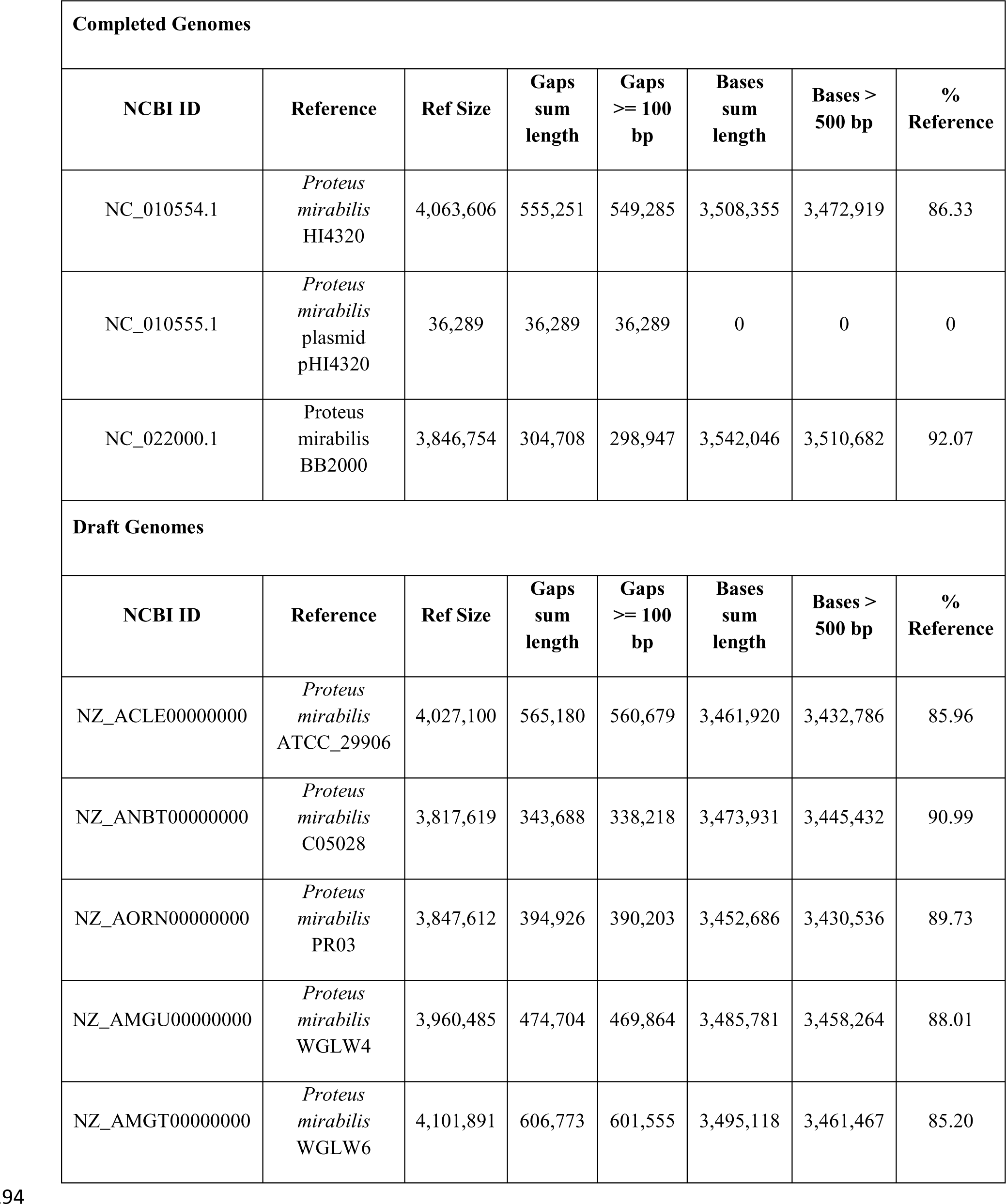
Comparison *of Proteus mirabilis* SCDR1 to complete and draft reference genomes *of Proteus mirabilis.*

**Figure 2:**
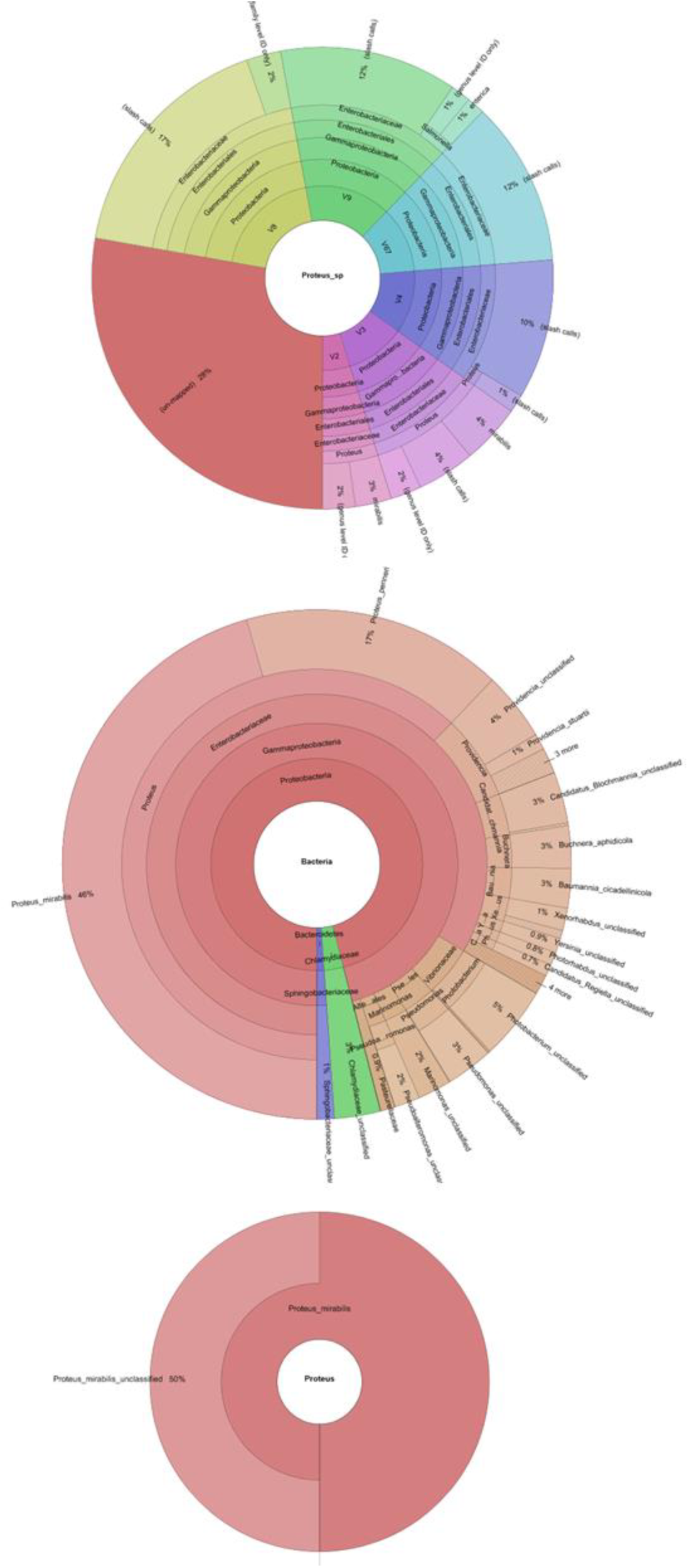
Metaphlan primary identification of the tested taxon.

A similar scenario was observed when constructing the phylogenetic relationship between our isolate and other *Proteus mirabilis* available in the NCBI- GenBank. 16Sr DNA-based maximum likelihood phylogenetic tree (Figure 3) showed that our isolate is located within a large clade that contains the majority *of Proteus mirabilis* strains and isolates. In addition, *P. mirabilis* SCDR1 showed to be closely related to the reference strain *P. mirabilis* HI4320 compared with *P. mirabilis* BB2000 that is located in another clade of four Proteus *mirabilis* taxa. On the contrary, whole genome Neighbor-joining phylogenetic tree of *Proteus mirabilis* spices including *P. mirabilis* SCDR1 isolate (Figure 4), showed that our isolate is more closely related to *P. mirabilis* BB2000 compared with the reference strain/genome *P. mirabilis* HI4320. However, Figure 4 showed that *P. mirabilis* SCDR1 exhibited obvious genetic divergence from other *Proteus mirabilis* species. Similar results were observed when performing pairwise pair-wise whole genome alignment of *P. mirabilis* strain SCDR1 against reference genomes (Figure 5).

**Figure 3:**
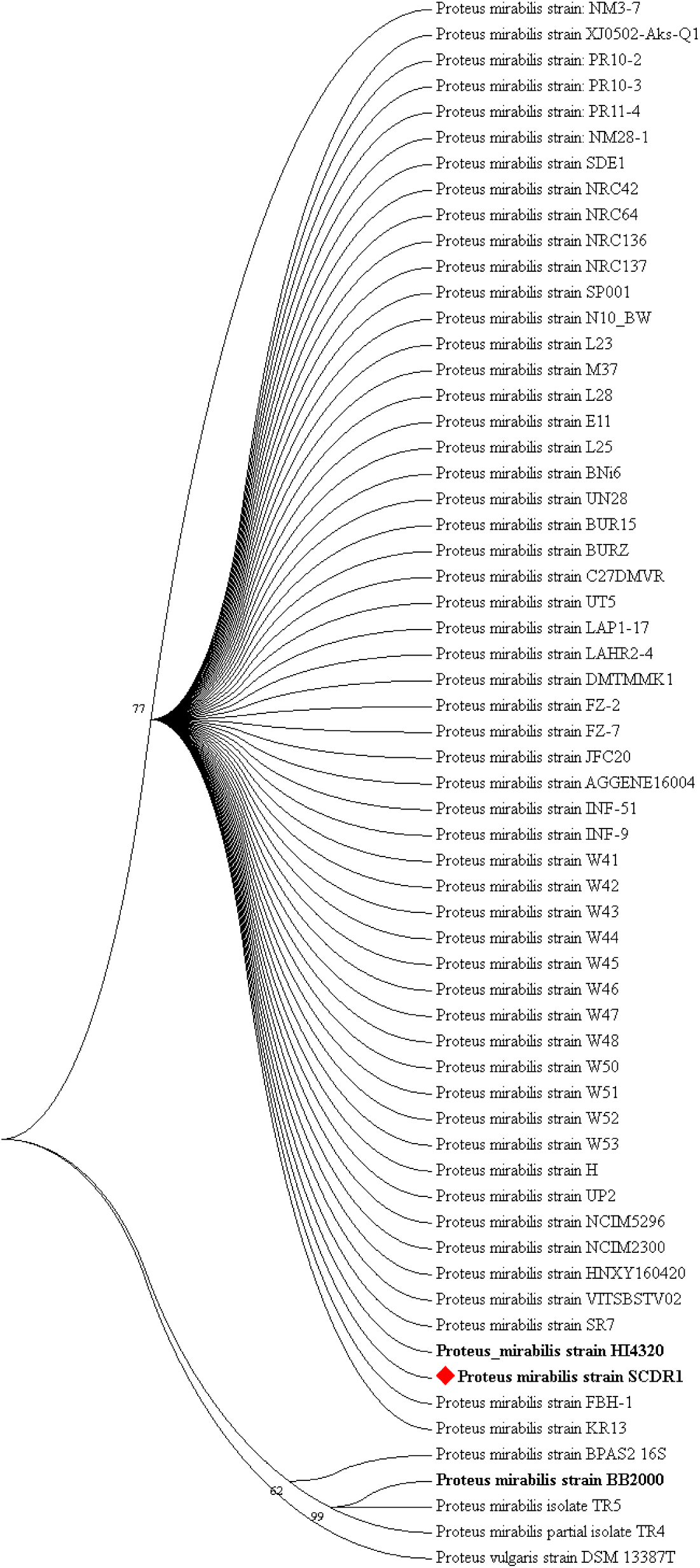
16S rDNA based Maximum likelihood phylogenetic tree of Proteus mirabilis spices including Pm-SCDR1 isolate.

**Figure 4:**
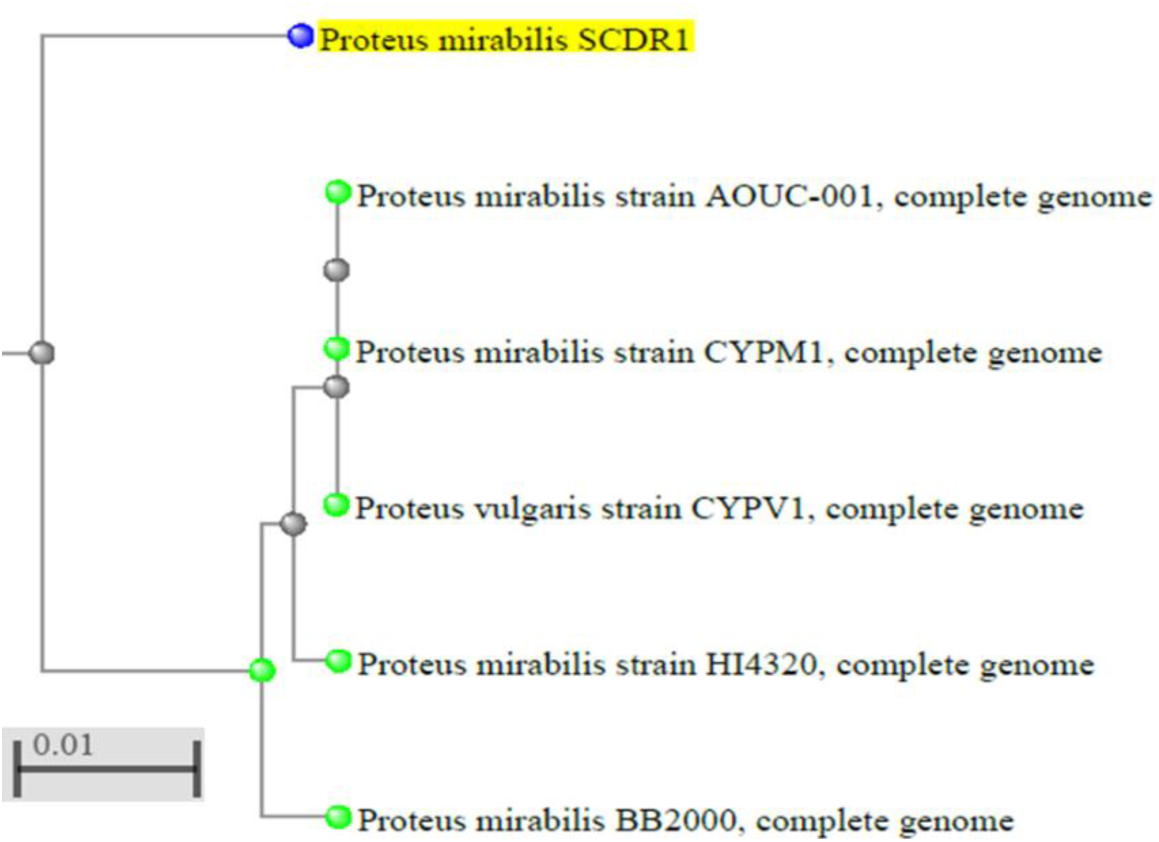
Whole genome Neighbor joining phylogenetic tree of *Proteus mirabilis* spices including Pm-SCDR1 isolate.

**Figure 5:**
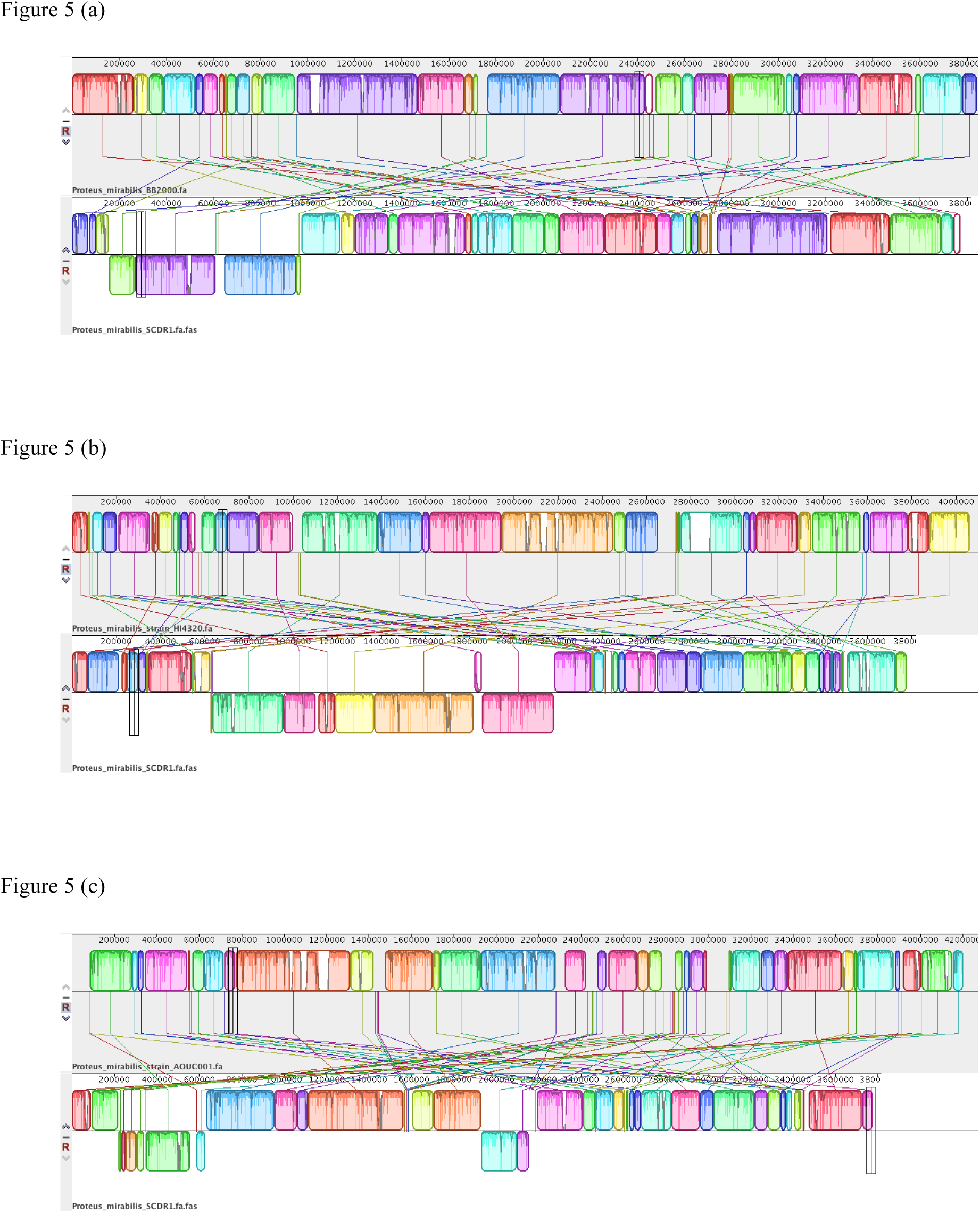
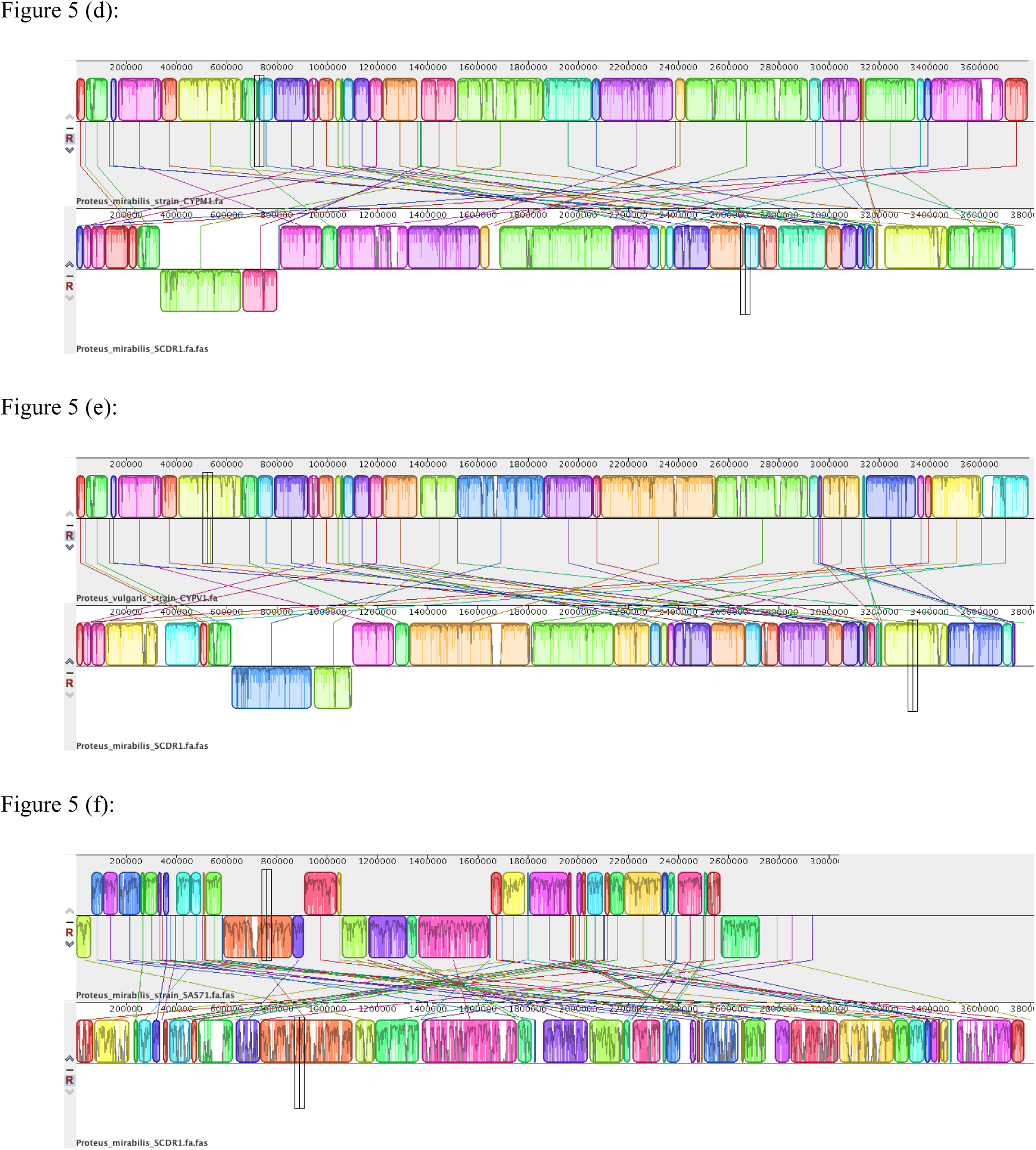
Pair-wise Whole Genome Alignment *of *P. mirabilis** strain SCDR1 against reference genomes. (a) *P. mirabilis BB200* and *P. mirabilis* SCDR1, (b) *P. mirabilis HI4320* and *P. mirabilis* SCDR1, (c) *P. mirabilis* AOUC001 and *P. mirabilis* SCDR1, (d) *P. mirabilis* CYPM1 and *P. mirabilis* SCDR1, (e) *P. vulgaris* CYPV1 and *P. mirabilis* SCDR1 and (f) *P. mirabilis* SAS71 and *P. mirabilis* SCDR1 Mauve whole genome alignment.

This was also confirmed with the clear divergence among *P. mirabilis* SCDR1 *Proteus mirabilis* species on the proteomic level (Figure 6). Comparing annotated proteins across genomes showed that the majority of protein sequence identity ranged between 95-99.5% with the highest values (100%) was observed for ribosomal proteins such as, SSU ribosomal protein S10p (S20e), LSU ribosomal protein L3p (L3e), LSU ribosomal protein L4p (L1e), and energy production involved proteins such as, ATP synthase gamma chain, beta chain and epsilon chain, cell division proteins such as, Cell division protein FtsZ, FtsA and FtsQ, NADH-ubiquinone oxidoreductase chains K, J, I, H and G and some other conserved essential proteins. On the other hand low values of protein identity similarities (26-85%) were observed for some proteins such as Fimbriae related proteins, transcriptional regulators, Ribosomal large subunit pseudouridine synthases, Phage-related proteins, O-antigen acetylases, inner and outer membrane-related proteins, secreted proteins, heavy metal transporting ATPases, Drug resistance efflux proteins, Iron transport proteins and cell invasion proteins.

**Figure 6:**
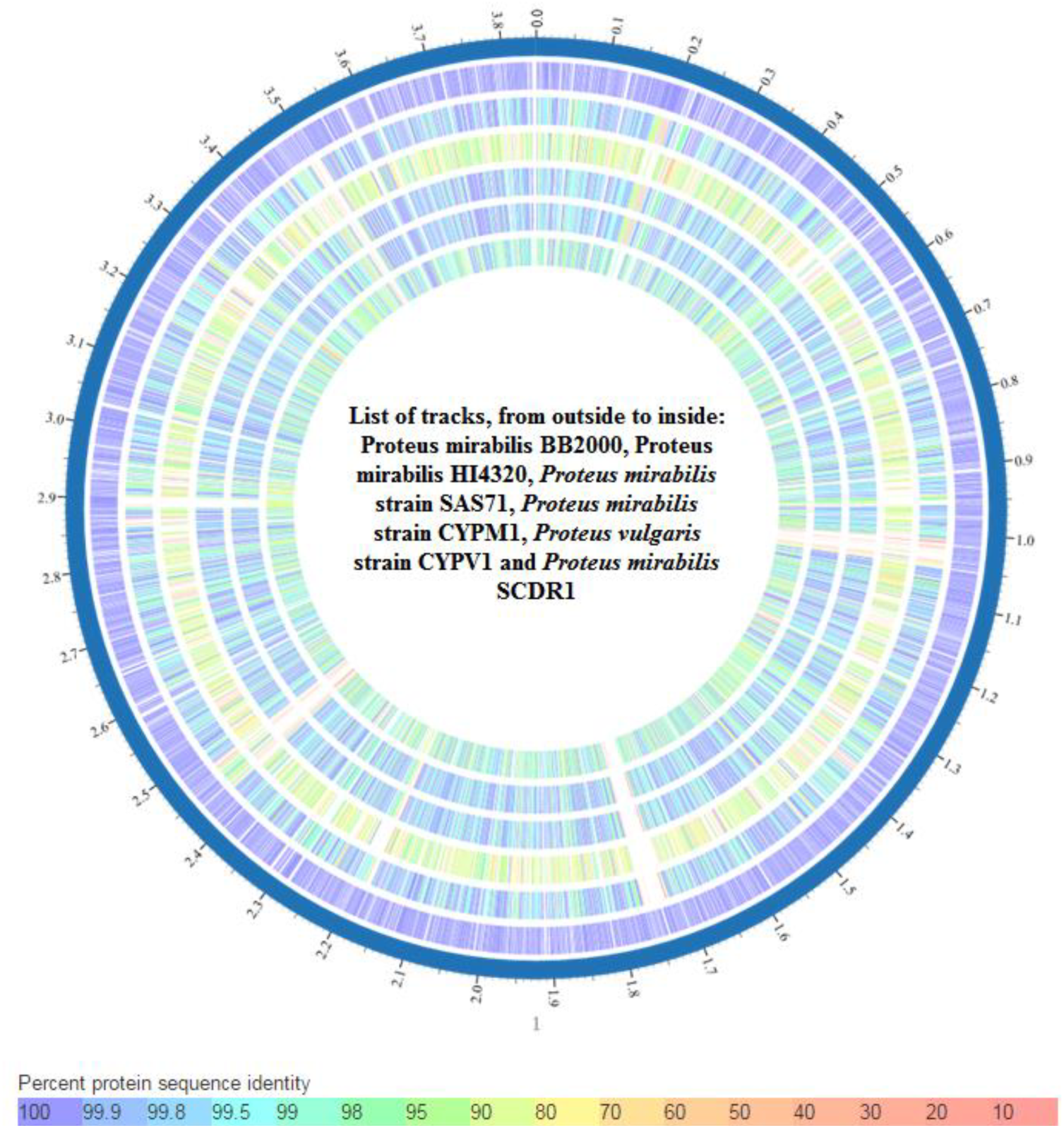
Whole genome phylogeny based proteomic comparison among *Proteus mirabilis* strains.

### Bacterial pathogenic and virulence factors

Pathogenomics analysis using PathogenFinder 1.1 showed that our input organism was predicted as a human pathogen, Probability of being a human pathogen 0.857. *P. mirabilis* SCDR1 comparative proteome analysis showed 35 matched hits from pathogenic families and only one hit from non-pathogenic families (**Supplementary Table 3**). In addition, genome analysis showed that *P. mirabilis* SCDR1 isolate contains numerous virulence factor genes and/or operons that marques it to be a virulent pathogenic bacterium. These virulence factors include Swarming behavior, mobility (flagellae), adherence, toxin and hemolysin production, Urease, Quorum sensing, iron acquisition systems, proteins that function in immune evasion, cell invasion and biofilm formation, stress tolerance factors, and chemotaxis related factors (**Supplementary Table 4**).

### Proteus mirabilis SCDR1 Resistome

#### Antibiotic resistance

Antibiotic resistance identification Perfect and Strict analysis using Resistance Gene Identifier (RGI) showed that *P. mirabilis* SCDR1 isolate contains 34 antibiotic resistance genes that serve in 21 antibiotic resistance functional categories (**Supplementary Table 5 and** Figure 7). Table 5 displays the consensus *P. mirabilis-SCDR1* antibiotic resistome. Genomics analysis of *P. mirabilis*-SCDR1 63 contigs showed that our isolates contains genetic determinants for tetracycline resistance (tetAJ), fluoroquinolones resistance (gyrA, parC and parE), sulfonamide resistance (folP), daptomycin and rifamycin resistance (rpoB), elfamycin antibiotics resistance (tufB), Chloramphenicol (cpxR, cpxA and cat), Ethidium bromide-methyl viologen resistance protein (emrE) and Polymyxin resistance (phoP). In addition, several multidrug resistance efflux systems and complexes such as MdtABC-TolC, MacAB-TolC, AcrAB-TolC, EmrAB-TolC, AcrEF-TolC and MATE.

**Figure 7:**
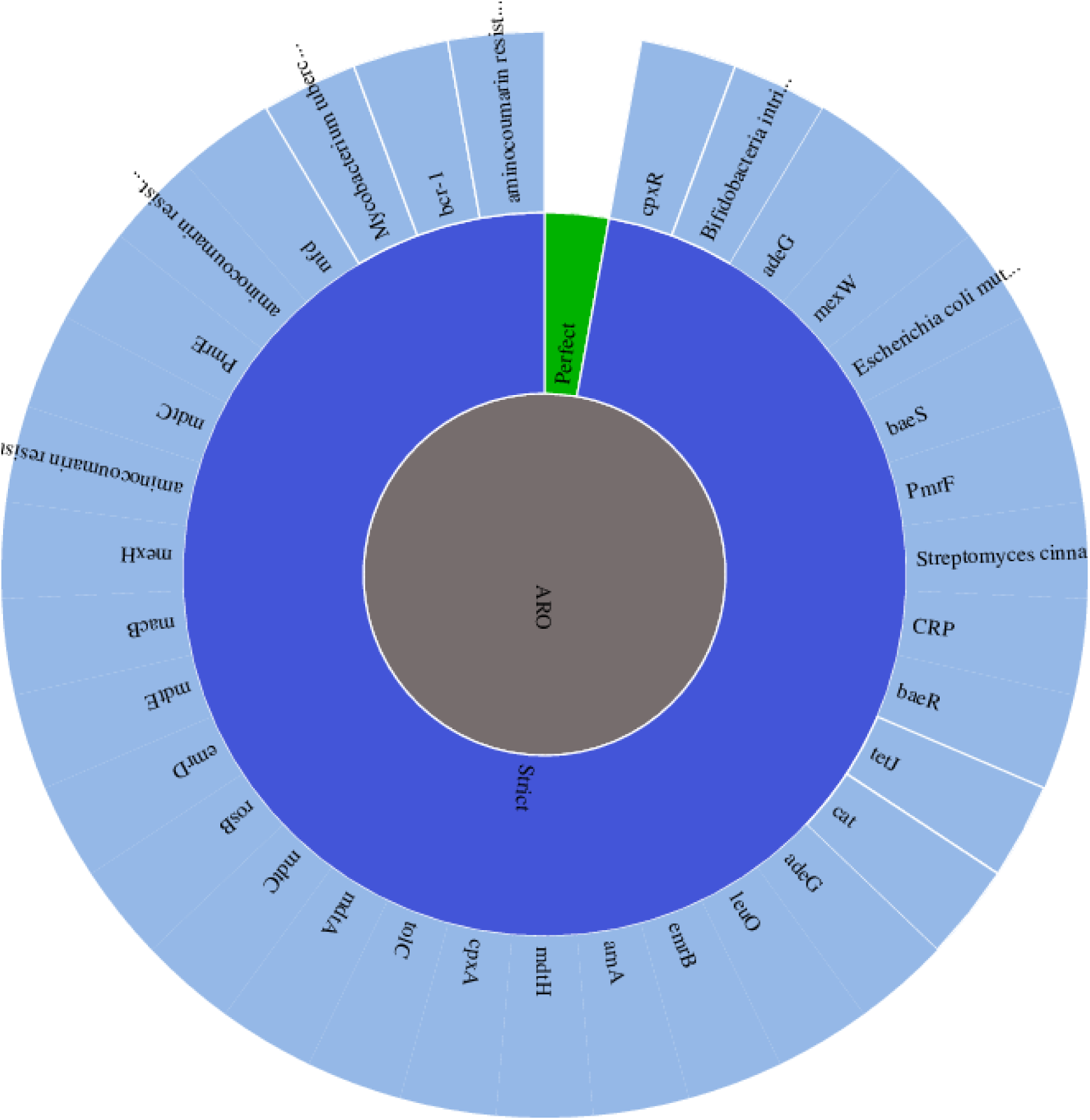
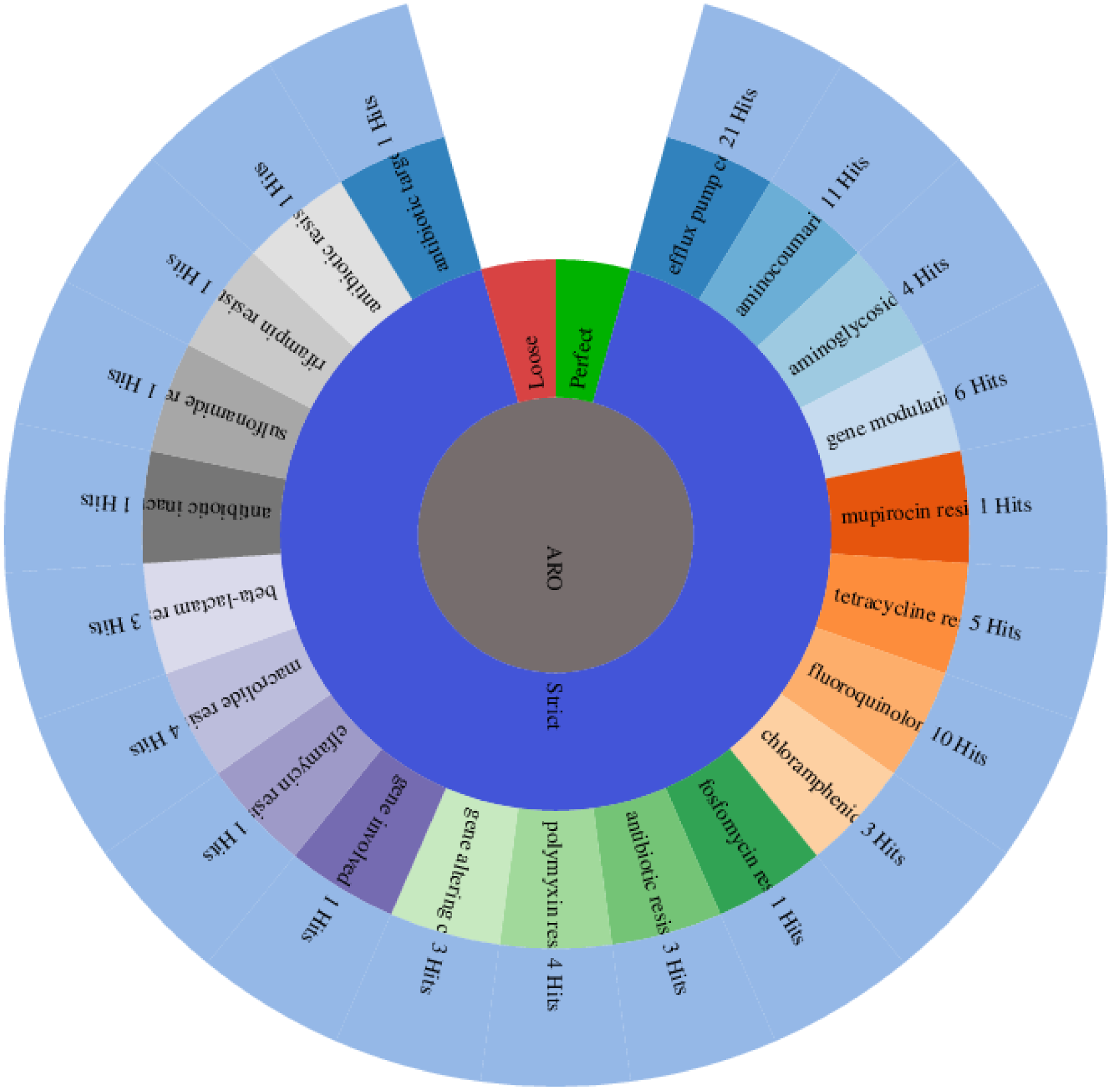
Antibiotic Resistance strict gene analysis and function analysis for Proteus mirabilis SCDR1.

**Table 5:** Consensus *P. mirabilis-SCDR1* antibiotic Resistome.

| Source | Source Organism | Gene | Product | Function | Query Coverage | Identity | E-value |
| --- | --- | --- | --- | --- | --- | --- | --- |
| ARDB | *P. mirabilis* ATCC 29906 | tetAJ | Tetracycline efflux protein TetA | Major facilitator superfamily transporter, tetracycline efflux pump. | 97 | 95 | 0 |
| CARD | *P. mirabilis* BB2000 | tetAJ | Tetracycline efflux protein TetA | Major facilitator superfamily transporter, tetracycline efflux pump. | 97 | 94 | 0 |
| ARDB | *P. mirabilis* HI4320 | tetAJ | Tetracycline efflux protein TetA | Major facilitator superfamily transporter, tetracycline efflux pump. | 80 | 99 | 2e-74 |
| CARD | *P. mirabilis* BB2000 | gyrA | DNA gyrase subunit A (EC 5.99.1.3) | Point mutation of *Escherichia coli* gyrA resulted in the lowered affinity between fluoroquinolones and gyrA. Thus, conferring resistance | 98 | 99 | 0 |
| CARD | *P. mirabilis* BB2000 | baeR | Response regulator BaeR | BaeR is a response regulator that promotes the expression of MdtABC and AcrD efflux complexes. | 100 | 99 | 2e-171 |
| CARD | *P. mirabilis* BB2000 | baeS | Sensory histidine kinase BaeS | BaeS is a sensor kinase in the BaeSR regulatory system. While it phosphorylates BaeR to increase its activity. | 100 | 99 | 0 |
| CARD | *P. mirabilis* BB2000 | mdtC | Multidrug transporter MdtC | MdtC is a transporter that forms a heteromultimer complex with MdtB to form a multidrug transporter. MdtBC is part of the MdtABC-TolC efflux complex. | 100 | 99 | 0 |
| CARD | *P. mirabilis* BB2000 | mdtB | Multidrug transporter MdtB | MdtB is a transporter that forms a heteromultimer complex with MdtC to form a multidrug transporter. MdtBC is part of the MdtABC-TolC efflux complex. | 100 | 99 | 0 |
| CARD | *P. mirabilis* BB2000 | mdtA | RND efflux system, membrane fusion protein | MdtA is the membrane fusion protein of the multidrug efflux complex mdtABC. | 100 | 98 | 0 |
| CARD | *P. mirabilis* BB2000 | folP | Dihydropteroate synthase (EC 2.5.1.15) | Point mutations in dihydropteroate synthase folP prevent sulfonamide antibiotics from inhibiting its role in folate synthesis, thus conferring sulfonamide resistance. | 100 | 100 | 0 |
| CARD | *P. mirabilis* BB2000 | soxR | Redox-sensitive transcriptional activator SoxR | SoxR is a sensory protein that upregulates soxS expression in the presence of redox-cycling drugs. This stress response leads to the expression many multidrug efflux pumps. | 100 | 100 | 0 |
| CARD | *Shigella dysenteriae* Sd197 | ompR | Two-component system response regulator OmpR | Transcriptional regulatory protein | 99 | 87 | 0 |
| CARD | *P. mirabilis* BB2000 | emrR | Transcriptional repressor MprA | EmrR is a negative regulator for the EmrAB-TolC multidrug efflux pump in *E. coli*. Mutations lead to EmrAB-TolC overexpression. | 100 | 100 | 0 |
| CARD | *P. mirabilis* BB2000 | emrA | Multidrug resistance protein ErmA | EmrA is a membrane fusion protein, providing an efflux pathway with EmrB and TolC between the inner and outer membranes of *E. coli*, a Gram-negative bacterium. | 95 | 96 | 0 |
| CARD | *P. mirabilis* BB2000 | acrE | Membrane fusion component of tripartite multidrug resistance system | AcrEF-TolC is a tripartite multidrug efflux system similar to AcrAB-TolC and found in Gram-negative bacteria. AcrE is the membrane fusion protein, AcrF is the inner membrane transporter, and TolC is the outer membrane channel protein. | 100 | 98 | 3e-44 |
| CARD | *P. mirabilis* BB2000 | emrB | Multidrug resistance protein ErmB | emrB is a translocase in the emrB-TolC efflux protein in *E. coli*. It recognizes substrates including carbonyl cyanide m-chlorophenylhydrazone (CCCP), nalidixic acid, and thioacetamycin. | 100 | 99 | 0 |
| CARD | *P. mirabilis* BB2000 | rpoB | DNA-directed RNA polymerase beta subunit (EC 2.7.7.6) | Mutations in rpoB gene confers antibiotic resistance (Daptomycin and Rifamycin) | 100 | 99 | 0 |
| CARD | *P. mirabilis*<br>BB2000 | tufB | Translation elongation factor Tu | Sequence variants of elongation factor Tu confer resistance to elfamycin antibiotics. | 100 | 100 | 1e-43 |
| CARD | *P. mirabilis*<br>BB2000 | cpxA | Copper sensory histidine kinase CpxA | cpxA mutant confer resistant to amikacin | 94 | 99 | 0 |
| CARD | *P. mirabilis*<br>BB2000 | cpxR | Copper-sensing two-component system response regulator CpxR | CpxR is a regulator that promotes acrD expression when phosphorylated by a cascade involving CpxA, a sensor kinase. cefepime and chloramphenicol | 100 | 100 | 0 |
| CARD | *P. mirabilis*<br>BB2000 | emrD | Multidrug resistance protein D | EmrD is a multidrug transporter from the Major Facilitator Superfamily (MFS) primarily found in Escherichia coli. EmrD couples efflux of amphipathic compounds with proton import across the plasma membrane. | 100 | 99 | 0 |
| CARD | *P. mirabilis*<br>BB2000 | macA | Macrolide-specific efflux protein MacA | MacA is a membrane fusion protein that forms an antibiotic efflux complex with MacB and TolC. | 100 | 99 | 3e-177 |
| CARD | *P. mirabilis*<br>BB2000 | macB | Macrolide export ATP-binding/permease protein MacB (EC 3.6.3.-) | MacB is an ATP-binding cassette (ABC) transporter that exports macrolides with 14- or 15- membered lactones. It forms an antibiotic efflux complex with MacA and TolC. | 100 | 98 | 0 |
| ARDB | *P. mirabilis*<br>ATCC 29906 | cat | Chloramphenicol acetyltransferase (EC 2.3.1.28) | Group A chloramphenicol acetyltransferase, which can inactivate chloramphenicol. | 99 | 93 | 6e-150 |
| CARD | *P. mirabilis*<br>BB2000 | cat | Chloramphenicol acetyltransferase (EC 2.3.1.28) | Group A chloramphenicol acetyltransferase, which can inactivate chloramphenicol. | 99 | 93 | 4e-151 |
| CARD | *P. mirabilis*<br>BB2000 | acrR | Transcription repressor of multidrug efflux pump acrAB operon, TetR (AcrR) family | AcrR is a repressor of the AcrAB-TolC multidrug efflux complex. AcrR mutations result in high level antibiotic resistance. | 100 | 95 | 9e-25 |
| CARD | *P. mirabilis*<br>BB2000 | acrR | Transcriptional regulator of acrAB operon, AcrR | AcrR is a repressor of the AcrAB-TolC multidrug efflux complex. AcrR mutations result in high level antibiotic resistance. | 93 | 95 | 2e-114 |
| CARD | *P. mirabilis*<br>BB2000 | acrA | RND efflux system, membrane fusion protein | Protein subunit of AcrA-AcrB-TolC multidrug efflux complex. AcrA represents the periplasmic portion of the transport protein. | 100 | 99 | 0 |
| CARD | *P. mirabilis*<br>BB2000 | mdtK | Multi antimicrobial extrusion protein (Na(+)/drug antiporter), MATE family of MDR efflux pumps | A multidrug and toxic compound extrusions (MATE) transporter conferring resistance to norfloxacin, doxorubicin and acriflavine. | 98 | 99 | 3e-164 |
| CARD | *Salmonella enterica*<br>subsp. *enterica*<br>serovar Agona str. SL483 | hns | DNA-binding protein H-NS | H-NS is a histone-like protein involved in global gene regulation in Gram-negative bacteria. It is a repressor of the membrane fusion protein genes acrE, mdtE, and emrK as well as nearby genes of many RND-type multidrug exporters. | 100 | 80 | 0 |
| CARD | *P. mirabilis*<br>BB2000 | tufB | Translation elongation factor Tu | Sequence variants of elongation factor Tu confer resistance to elfamycin antibiotics. | 100 | 99 | 0 |
| CARD | *Shigella dysenteriae*<br>Sd197 | crp | Cyclic AMP receptor protein | CRP is a global regulator that represses MdtEF multidrug efflux pump expression. | 100 | 98 | 0 |
| CARD | *P. mirabilis*<br>BB2000 | emrE | Ethidium bromide-methyl viologen resistance protein EmrE | EmrE is a small multidrug transporter that functions as a homodimer and that couples the efflux of small polyaromatic cations from the cell with the import of protons down an electrochemical gradient. EmrE is found in E. coli and P. aeruginosa. | 100 | 99 | 6e-73 |
| CARD | *P. mirabilis*<br>BB2000 | mdtK | Multi antimicrobial extrusion protein (Na(+)/drug antiporter), MATE family of MDR efflux pumps | A multidrug and toxic compound extrusions (MATE) transporter conferring resistance to norfloxacin, doxorubicin and acriflavine. | 100 | 100 | 2e-113 |
| CARD | *P. mirabilis*<br>BB2000 | NIA | Putative transport protein | NIA | 100 | 94 | 7e-59 |
| CARD | *P. mirabilis*<br>BB2000 | NIA | Multidrug resistance protein | NIA | 99 | 96 | 2e-112 |
| CARD | *P. mirabilis*<br>BB2000 | parC | Topoisomerase IV subunit A (EC 5.99.1.-) | ParC is a subunit of topoisomerase IV, which decatenates and relaxes DNA to allow access to genes for transcription or translation. Point mutations in ParC prevent fluoroquinolone antibiotics from inhibiting DNA synthesis, and confer low-level resistance. Higher-level resistance results from both gyrA and parC mutations. | 99 | 99 | 0 |
| CARD | *P. mirabilis*<br>BB2000 | parE | Topoisomerase IV subunit B (EC 5.99.1.-) | ParE is a subunit of topoisomerase IV, necessary for cell survival. Point mutations in ParE prevent fluoroquinolones from inhibiting DNA synthesis, thus conferring resistance. | 100 | 99 | 0 |
| CARD | *P. mirabilis*<br>BB2000 | tolC | Type I secretion outer membrane protein, TolC precursor | TolC is a protein subunit of many multidrug efflux complexes in Gram negative bacteria. It is an outer membrane efflux protein and is constitutively open. Regulation of efflux activity is often at its periplasmic entrance by other components of the efflux complex. | 100 | 99 | 0 |
| CARD | *P. mirabilis*<br>BB2000 | mdtH | MFS superfamily export protein YceL | Multidrug resistance protein MdtH | 100 | 99 | 0 |
| CARD | *P. mirabilis*<br>BB2000 | phoP | Transcriptional regulatory protein PhoP | A mutant phoP activates pmrHFIJKLM expression responsible for L-aminoarabinose synthesis and polymyxin resistance, by way of alteration of negative charge | 100 | 99 | 5e-165 |
| CARD | *P. mirabilis*<br>BB2000 | phoQ | Sensor histidine kinase PhoQ (EC 2.7.13.3) | Mutations in *Pseudomonas aeruginosa* PhoQ of the two-component PhoPQ regulatory system. Presence of mutation confers resistance to colistin | 90 | 99 | 0 |
| CARD | *P. mirabilis*<br>BB2000 | phoQ | Sensor histidine kinase PhoQ (EC 2.7.13.3) | Mutations in *Pseudomonas aeruginosa* PhoQ of the two-component PhoPQ regulatory system. Presence of mutation confers resistance to colistin | 98 | 98 | 1e-45 |

**Evidence:** BLASTP, **NIA:** No information available, **ARDB:** Antibiotic Resistance Genes Database, **CARD:** Comprehensive Antibiotic Resistance Database.

#### Heavy metal resistance

Table 6 presents *P. mirabilis* SCDR1 Heavy Metal Resistance/Binding factors. Numerous genetic determinants for metal resistance were observed in *P. mirabilis* SCDR1 genome. Several Copper resistance genes/proteins were detected, namely, copA, copB, copC, copD, cueO, cueR, cutC, cutF and CuRO_2_CopA_like1. In addition, gene determinants of Copper/silver efflux system were also observed, namely, oprB, oprM and cusC_1. Moreover, several heavy metal resistance proteins and efflux systems were also observed such as magnesium/cobalt efflux protein CorC, metal resistance proteins (AGS59089.1, AGS59090.1 and AGS59091.1), nickel-cobalt-cadmium resistance protein NccB, arsenical pump membrane protein (ArsB permease), Lead, cadmium, zinc and mercury transporting ATPase, outer membrane component of tripartite multidrug resistance system (CusC) and complete *P. mirabilis* tellurite resistance loci (terB, terA, terC, terD, terE, terZ). Furthermore, enzymes involved in heavy metal resistance were also observed such as Glutathione S-transferase (gst1, gst, Delta and Uncharacterized), arsenite S-371 adenosylmethyltransferase (Methyltransferase type 11) and alkylmercury lyase (MerB).

**Table 6:**
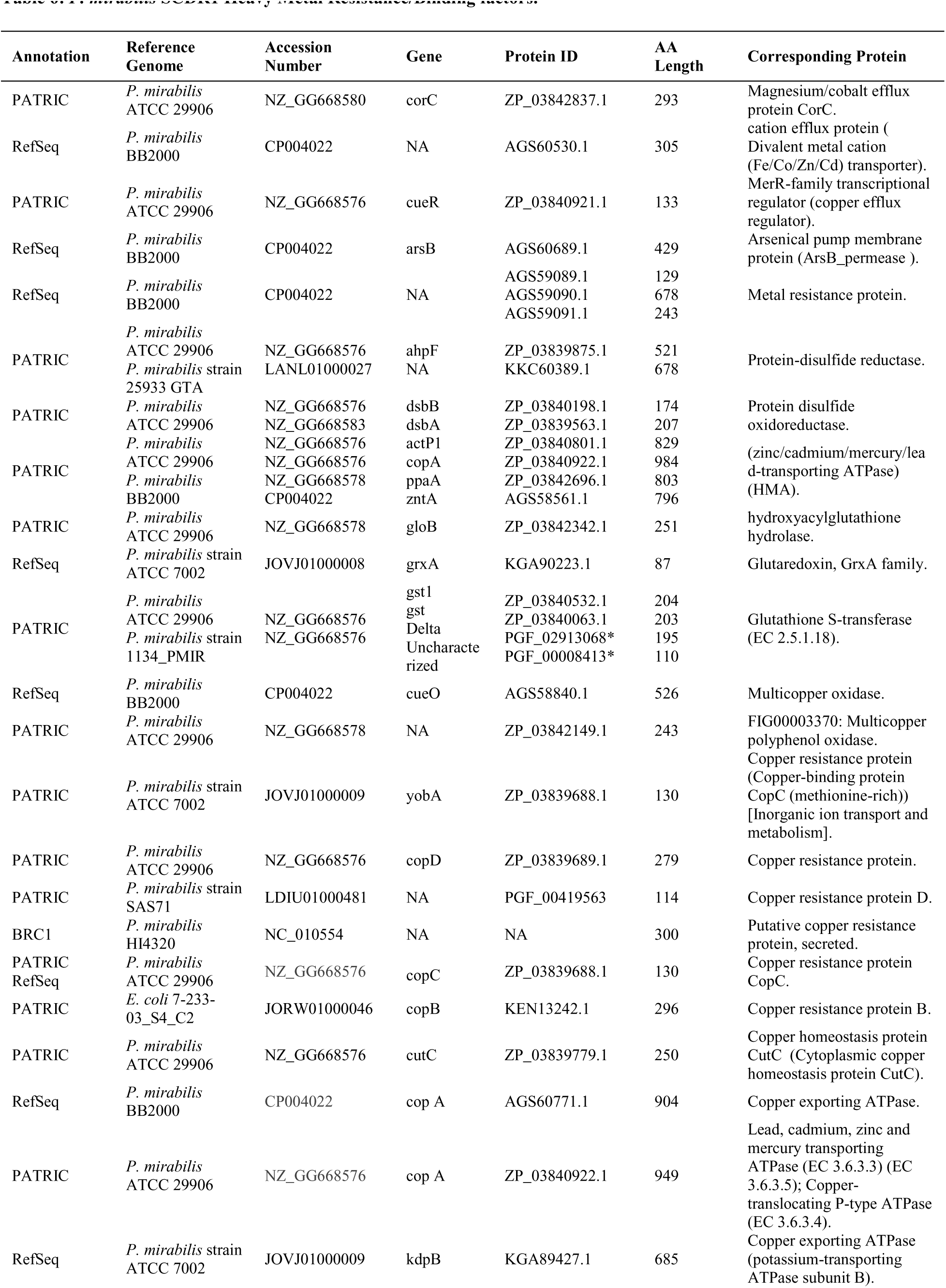
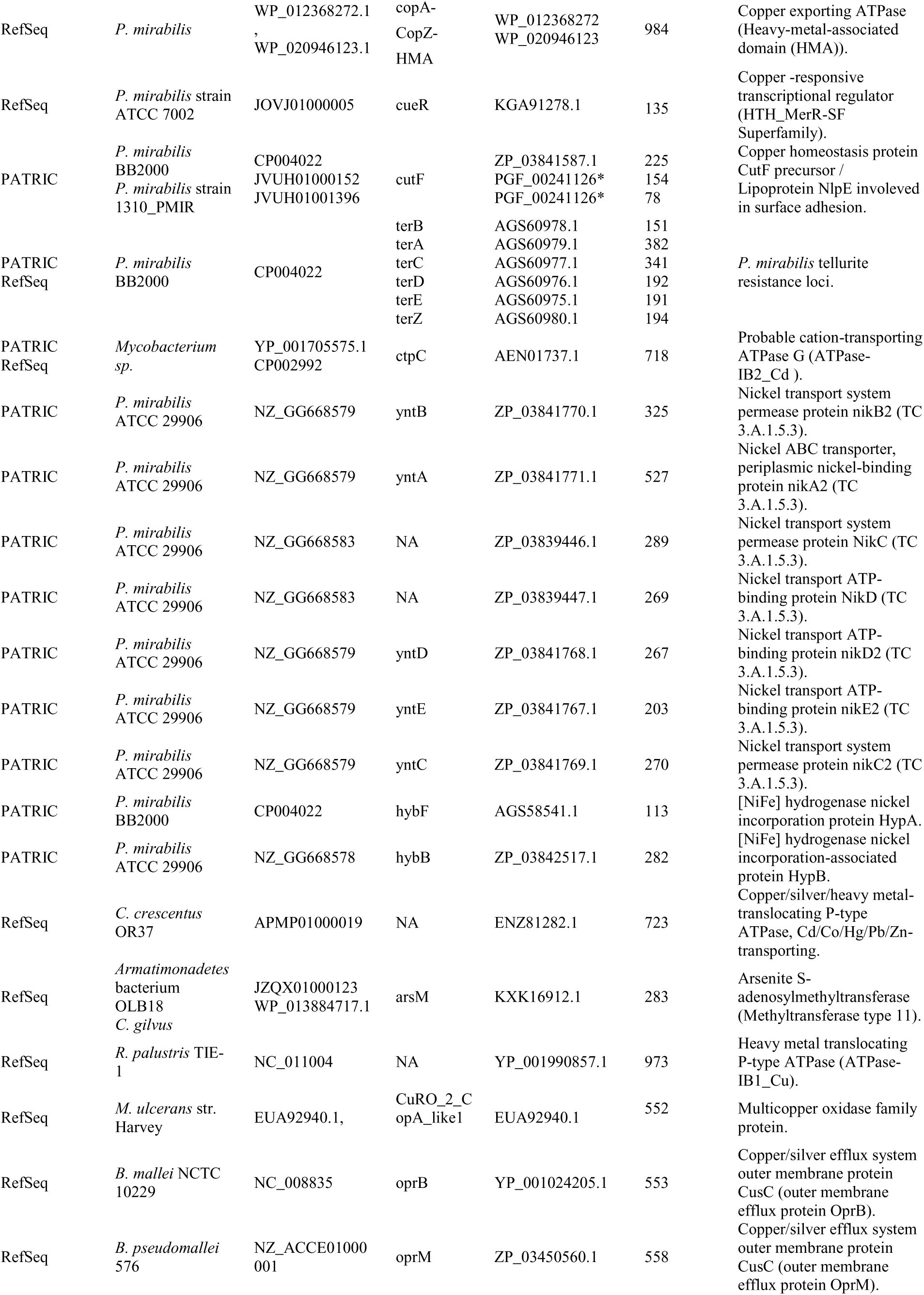
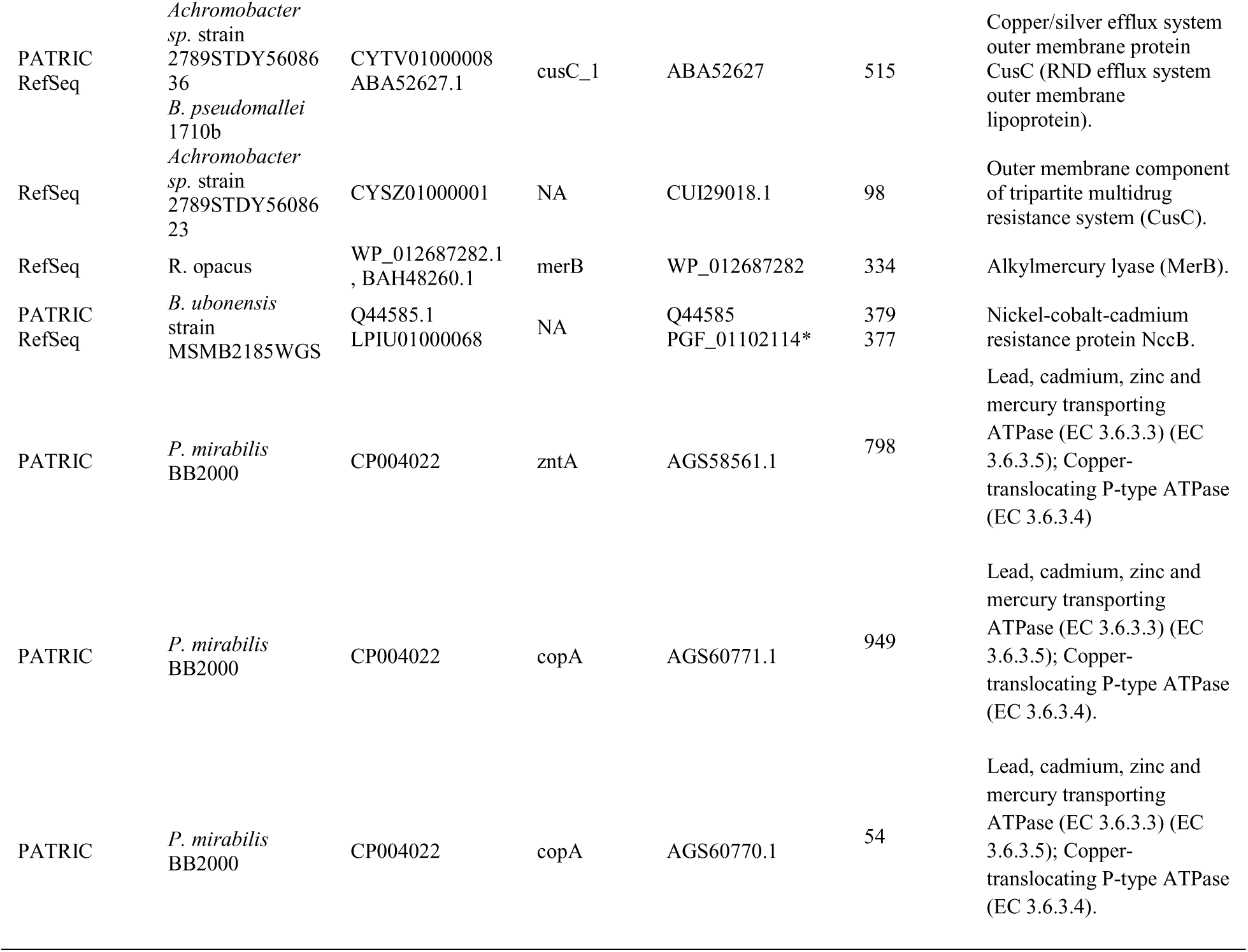
*P. mirabilis* SCDR1 Heavy Metal Resistance/Binding factors.

## Discussion

*Proteus mirabilis* SCDR1 isolate was isolated from a Diabetic ulcer patient visiting the Diabetic foot unit unit in the University Diabetes Center at King Saud University in the University Diabetes Center at King Saud University. Our SCDR1 isolate was observed as mixed culture along with *S. aureus* isolate while testing our produced silver Nanoparticles against several pathogenic *S. aureus* isolates (Saeb et al. 2014). Whereas other tested Gram positive and negative bacteria showed great sensitivity against silver Nanoparticles, *P. mirabilis* SCDR1 isolate exhibited extreme resistance. *P. mirabilis* SCDR1 isolate is multi-drug resistant bacteria (MDR), since, our isolate was non-susceptible to at least one agent in at least three antimicrobial categories (Magiorakos et al. 2012). Our isolate was against ansamycins, glycopeptides, fucidanes, cyclic peptides, nitroimidazoles, macrolides, lincosamides, folate pathway inhibitors and aminocoumarin antimicrobial categories. Moreover, our isolate exhibited the intrinsic resistant against tetracyclines and polymyxins specific to *P. mirabilis* species (Chen et al. 2015). However, fortunately, our isolates is sensitive against several operational antimicrobial categories such as penicillins with b-lactamase inhibitors, extended-spectrum cephalosporins, carbapenems, aminoglycosides, fluoroquinolones and phosphonic acids. In addition, our *P. mirabilis* SCDR1 isolate showed high resistance against colloidal and composite Nanosilver and metallic silver when compared to other tested Gram positive and negative bacterial species both qualitatively and quantitatively. To our knowledge, this is the first reported case of bacterial spontaneous resistance to colloidal and composite Nano-Silver. **P. mirabilis* SCDR1* demonstrated resistance against colloidal Nanosilver assessed either by disk diffusion or by minimal inhibitory concentration methods. While, all used concentrations of colloidal Nanosilver have shown strong effects on all tested microorganisms (Table 1), no effect on the bacterial growth of **P. mirabilis* SCDR1* even at the highest used concentration (200 ppm). Similarly, **P. mirabilis* SCDR1* were able to resist ten folds (500 ppm) higher than *K. pneumoniae* (50 ppm), five folds higher than *P. aeruginosa* and *E. coli* (100 ppm) and two and a half folds (200 ppm) higher than *S. aureus* and *E. cloacae* (Table 2). Moreover, while both laboratories prepared and commercially available silver and Nanosilver composite showed a clear effect against both *S. aureus* and *P. aeruginosa* the most common pathogens of diabetic foot ulcer, not effect was observed against **P. mirabilis* SCDR1* (Table 3). Although chitosan nanosilver composites have documented combined effect against both Gram positive and negative pathogens (Latif et al. 2015) no effect was observed against **P. mirabilis* SCDR1.*

**P. mirabilis* SCDR1* genome analysis showed that our isolate contains a large number of genes (245) responsible for xenobiotics biodegradation and metabolism (**supplementary table 2**). These includes Atrazine, Naphthalene and Trinitrotoluene degradation. In addition, we detected the presence genes encoding for the members Chitosanase family GH3 of N, N'-diacetylchitobiose-specific 6-phospho-beta-glucosidase (EC 3.2.1.86), Beta N-acetyl-glucosaminidase (nagZ, beta-hexosaminidase) (EC 3.2.1.52), and Glucan endo-1, 4-beta-glucosidase (EC 3.2.1.-) in *P. mirabilis* SCDR1 suggests that it can hydrolyze chitosan to glucosamine (Wieczorek et al. 2014; Gupta et al. 2010, 2011). This justifies the lack of antimicrobial effect of chitosan against **P. mirabilis* SCDR1.*

Similarly, *P. mirabilis* SCDR1 showed resistance against all the tested commercially available silver and Nanosilver containing wound dressing bandages. These silver containing commercially available bandages (wound dressing material) use different manufacturing technology and constituents. For example, Silvercel wound dressing contains high G calcium alginate in addition to 28% Silver-coated fibers (dressing contains 111mg silver/100cm^2^). The silver-coated fibers encompass elemental silver, which is converted to silver oxide upon contact with oxygen. Silver oxide dissolves in fluid and releases ionic silver (Ag^+^) that have antimicrobial action (Cutting et al. 2007). Clinical studies showed that Silvercel wound dressing is effective against many common wound pathogens, including methicillin-resistant Staphylococcus aureus (MRSA), methicillin -resistant Staphylococcus epidermidis (MRSE) and vancomycin-resistant Enterococcus (VRE). In addition, these studies showed that Silvercel wound dressing prevented and disrupted the formation of bacterial biofilms (McInroy et al. 2010; Stephens et al. 2010). However, this was not the case with our *P. mirabilis* SCDR1 isolate.

Pathogenomics analysis showed that *P. mirabilis* SCDR1 isolate is a potential virulent pathogen that despite its original isolation site, wound, it can establish kidney infection and its associated complications (**Supplementary tables 3 and 4**). *P. mirabilis* SCDR1 showed that it possesses the characteristic bull’s eye pattern swarming behavior. Presenting swarmer cells form is associated with the increase of expression of virulence genes (Allison et al. 1992). Swarming is important to *P. mirabilis* uropathogenesis. When this microorganism presents swarmer cells form, the expression of virulence is increased (Allison et al. 1992). It was shown that swarming bacteria that move in multicellular groups exhibit adaptive resistance to multiple antibiotics (Butler et al. 2010). Moreover, migrating swarm cells display an increased resistance many of antimicrobial agents. For example, swarm cells of *P. aeruginosa* were able to migrate very close to the disc containing arsenite, indicating resistance to this heavy metal (Lai et al. 2009). It was suggested that high densities promote bacterial survival, the ability to move, as well as the speed of movement, confers an added advantage, making swarming an effective strategy for prevailing against antimicrobials including heavy metals (Lai et al. 2009; Butler et al. 2010). Altruism or self-sacrifice is a suggested phenomenon associated with swarming that involves risk of wiping out some individuals upon movement of bacteria to a different location allowing the remaining individuals to continue their quest (Butler et al. 2010; Gadagkar 1997). Thus maintaining high cell density, though the observed quorum sensing ability (**supplementary table 4**), circulating within the multilayered colony to minimize exposure to the heavy metal, and the death of individuals that are directly exposed could be the key to the observed Nanosilver resistance.

*P. mirabilis* SCDR1 isolate exhibited the ability of biofilm formation and also our pathogenomics analysis showed that it contains genes responsible for it such as glpC gene coding for anaerobic glycerol-3-phosphate dehydrogenase subunit C (EC 1.1.5.3), pmrI gene coding for UDP-glucuronic acid decarboxylase and baaS gene coding for biofilm formation regulatory protein BssS. Uropathogens use different mechanisms including biofilm formation for survival in response to stresses in the bladder such as starvation and immune responses (Justice et al. 2008; Horvath et al. 2011). Also, biofilm formation can reduce the metal toxic effect by trapping it outside the cells. It was found that in the relative bacteria *Proteus vulgaris* XC 2 the biofilm cells of showed considerably greater resistance to Chloromycetin compared to planktonic cells (free-floating counterparts) (Wu et al. 2015). In addition, it was found that biofilm formation and exopolysaccharide are very important for the heavy metal resistance in *Pseudomonas* sp. and that biofilm lacking mutant was less tolerant to heavy metals (Chien et al. 2013). Furthermore, it was found that both extracellular polysaccharides and biofilm formation is a resistance mechanism against to toxic metals in *Sinorhizobium meliloti,* the nitrogen-fixing bacterium (Nocelli et al. 2016). Thus, we suggest that the ability of *P. mirabilis* SCDR1 to form biofilm may also assist in the observed Nanosilver resistance.

In addition, *P. mirabilis* SCDR1 contains several genes and proteins that also facilitate metal resistance including silver and Nanosilver (table 6). We observed the presence of gene determinants of Copper/silver efflux system, oprB encoding for Copper/silver efflux system outer membrane protein CusC (outer membrane efflux protein OprB), oprM encoding for Copper/silver efflux system outer membrane protein CusC (outer membrane efflux protein OprM), cusC_1 encoding for Copper/silver efflux system outer membrane protein CusC (RND efflux system outer membrane lipoprotein), cpxA encoding for Copper sensory histidine kinase and outer membrane component of tripartite multidrug resistance system (CusC). Indicating the presence of endogenous silver and copper resistance mechanism in *P. mirabilis* SCDR1. Similar endogenous silver and copper resistance mechanism has been described in *E. coli* has been associated with loss of porins from the outer membrane and up-regulation of the native Cus efflux mechanism that is capable of transporting silver out of the cell (Li et al. 1997; Lok et al. 2008). Thus we suggest a comprehensive study for this endogenous silver resistance mechanism within *Proteus mirabilis* species.

Furthermore, we observed the presence of enzymes involved in heavy metal resistance such as Glutathione S-transferase (EC 2.5.1.18) (gst1, gst, Delta and Uncharacterized) in *P. mirabilis* SCDR1 genome. Glutathione S-transferases (GSTs) are a family of multifunctional proteins playing important roles in detoxification of harmful physiological and xenobiotic compounds in organisms (Zhang et al. 2013). Moreover, it was found that a Glutathione S-transferase is involved in copper, cadmium, lead and mercury resistance (Nair and Choi 2011). Furthermore, it was found that GST genes are differentially expressed in defense against oxidative stress caused by Cd and Nanosilver exposure (Nair and Choi 2011). Thus we can propose a role of Glutathione S-transferases of *P. mirabilis* SCDR1 in the observed Nanosilver resistance. Moreover, we observed the presence of a complete tellurite resistance operon (terB, terA, terC, terD, terE, terZ) that was suggested to contribute to virulence or fitness and protection from other forms of oxidative stress or agents causing membrane damage, such as silver and Nanosilver, in *P. mirabilis* (Toptchieva et al. 2003).

Several other heavy metal resistance genes and proteins were observed in the *P. mirabilis* SCDR1 genome. Such as, arsM encoding for arsenite S-adenosylmethyltransferase (Methyltransferase type 11) that play important role in prokaryotic resistance and detoxification mechanism to arsenite (Qin et al. 2009, 2006) and merB encoding for alkylmercury lyase that cleaves the carbon-mercury bond of organomercurials such as phenylmercuric acetate (Marchler-Bauer et al. 2015).

In addition, we observed the presence of several multidrug resistance efflux systems and complexes such as MdtABC-TolC, which is a multidrug efflux system in Gram-negative bacteria, including *E. coli* and *Salmonella* that confer resistance against β-lactams, novobiocin and deoxycholate (Nishino et al. 2007). It is noteworthy to mention that MdtABC-TolC and AcrD paly role in metal resistance (copper and zinc) along with their BaeSR regulatory system (Franke et al. 2003) that also was found in our *P. mirabilis* SCDR1 genome [table 5] thus also may play additional role in silver resistance. The MdtABC and AcrD systems may be related to bacterial metal homeostasis by transporting metals directly. This is to some extent similar to the copper and silver resistance mechanism by cation efflux of the CusABC system belonging to the RND protein superfamily (Franke et al. 2003; Outten et al. 2001). In addition, our isolate contains MacAB-TolC efflux pump which is an ABC efflux pump complex expressed in *E. coli* and *Salmonella enterica* and confers resistance to macrolides, including erythromycin (Nishino et al. 2006). Furthermore, we detected that presence of AcrAB-TolC efflux pump which is a tripartite RND efflux system that confers resistance to tetracycline, chloramphenicol, ampicillin, nalidixic acid, and rifampin in Gram-negative bacteria (Tikhonova et al. 2011). Moreover, EmrAB-TolC efflux system that confer resistance to nalidixic acid and thiolactomycin was also observed (Lomovskaya et al. 1995). In addition, AcrEF-TolC, which is a tripartite multidrug efflux system similar to AcrAB-TolC, was found in Gram-negative bacteria (Zheng et al. 2009). Finally, Multidrug and toxic compound extrusion (MATE) system was observed in *P. mirabilis-SCDR1* genome. It is responsible for Directed pumping of antibiotic out of a cell and thus of resistance. It utilizes the cationic gradient across the membrane as an energy source. Generally, the resistance gene search, resistome analysis, was in great agreement with the antibiotic sensitivity testing with very few exceptions. For example, several chloramphenicol resistance genes and proteins such as cpxR, cpxA, cat and AcrAB-TolC efflux pump were observed, though our *P. mirabilis* SCDR1 isolate was chloramphenicol sensitive. Yet genomic resistome analysis proofed to be a successful way of testing drug resistance and even discovering potential drug resistance genes in a given bacterium.

It is also worth mentioning that some cases we observed *P. mirabilis* SCDR1 adaptive resistance against and/secondary waves of swarming some antibiotics that initially scored as sensitive. These antibiotic belongs to the aminoglycosides (Spectinomycin and Streptomycin), cephalosporins (Ceftriaxone, Cefoxitin, Cephalothin, Cefotaxime, Cefaclor and Cefepime) and β-lactams (Aztreonam and Meropenem). Similar observations were also detected with *B. subtilis, B. thailandensis, E. coli* and (Lai et al. 2009) *Salmonella enterica* serovar Typhimurium (Tikhonova et al. 2011). Adaptation, rather than mutation, to increasing levels of antibiotics was suggested to justify the observed swarm waves.

The increasing antimicrobial nanosilver usage could prompt a silver resistance problem in Gram-negative pathogens, particularly since silver resistance is already known to exist in several such species (Li et al. 1997; Andersson 2003). Both exogenous (horizontally acquired Sil system) endogenous (mutational Cus system) resistance to silver has been reported in Gram-negative bacteria (Li et al. 1997; McHugh et al. 1975). Similarly, in our case we observed the presence of resistance operon with high similarity with the *cus* operon that is, in turn, is chromosomally encoded system because of the lack of any plasmid in *P. mirabilis* SCDR1. However, both endogenous and exogenous silver resistance systems, in Gram-negative bacteria, remain incompletely understood (Randall et al. 2015).

The occurrence of induced nanosilver resistance (in vitro) in *Bacillus sp.* (Gunawan et al. 2013), spontaneous resistance (in our case) and the frequent uses and misuses of nanosilver-containing medical products should suggest adopting an enhanced surveillance systems for nanosilver-resistant isolates in the medical setups. In addition, greater control over utilizing nanosilver-containing products should also be adapted in order to maintain nanosilver as valuable alternative approach for fighting multidrug resistant pathogens.

## Conclusion

In the present study, we introduced the **P. mirabilis* SCDR1* isolate that was collected from a Diabetic ulcer patient. **P. mirabilis* SCDR1* showed high levels of resistance against Nano-silver colloids, Nano-silver chitosan composite and the commercially available Nano-silver and silver bandages. Our isolate contains all the required pathogenicity and virulence factors to establish a successful infection. **P. mirabilis* SCDR1* contains several physical and biochemical mechanisms for antibiotics and silver/nanosilver resistance, which are biofilm formation, swarming mobility, efflux systems, and enzymatic detoxification.

## Acknowledgement

The authors want to thank the members of the Diabetic foot unit in the University Diabetes Center at King Saud University for their help in collecting the bacteria samples. Furthermore, we want to thanks the members of the nanotechnology department in SCDR for providing the chitosan nanosilver composites. In addition we want to acknowledge that NGS experiments and analysis were supported by the Saudi Human Genome Program (SHGP) at KACST and KFSHRC. Moreover, we want to thank Dr. Rebecca Wattam, form the Biocomplexity Institute at Virginia Polytechnic Institute and State University, for her great assistance during data analysis using PATRIC services and tools.

## List of abbreviations

**NGS:** Next generation sequencing techniques

**16S rRNA:** 16S ribosomal RNA gene

**Mb:** Mega base pairs

**GC content:** guanine-cytosine content

**BLASTn:** Basic Local Alignment Search Tool nucleotide

**bp:** Base pair

**SCDR:** Strategic center for Diabetes research

**KFSHRC:** King Faisal Specialist Hospital and Research Center

**PATRIC:** Pathosystems recourse Integration center

**DFU:** Diabetic foot ulcer

**MDR:** multidrug-resistant

**PPM:** part per million

**tRNAs:** Transfer ribonucleic acid

**AROs:** Antibiotic Resistance Ontology

**AMRO:** Antimicrobial Resistance based ontology

**RGI:** Resistance Gene Identifier

**DDT:** 1, 1, 1-Trichloro-2, 2-bis (4-chlorophenyl) ethane

**MRSA:** methicillin-resistant Staphylococcus aureus

**MRSE:** methicillin -resistant Staphylococcus epidermidis

**VRE:** Vancomycin-resistant Enterococcus

**MIC:** Minimum Inhibitory Concentration

**RND:** Resistance-Nodulation- Division

## Declarations

### * Ethics approval and consent to participate

This study was approved by institutional review board in King Saud University, Collage of Medicine Riyadh, Kingdom of Saudi Arabia. The subject was provided written informed consent for participating in this study.

### * Consent to publish

All other have consented for publication of this manuscript.

### * Availability of data and materials

Data from our draft genome of *P. mirabilis* SCDR1 isolate was deposited in NCBI-GenBank with an accession number LUFT00000000.

### * Authors’ contributions

**ATMS:** Involved in study conception and design, data analysis and interpretation. Involved in drafting the manuscript or revising it critically for important intellectual content. Preparing the final approval of the version to be published.

**KA:** Involved in study conception and design. Preparing the final approval of the version to be published.

**MAH:** Involved in study design. Involved in acquisition of data, or analysis and interpretation of data; preparation and involved in drafting the manuscript.

**MS:** Involved in acquisition of data, or analysis and interpretation of data.

**HT:** Involved in study conception and design. Involved in drafting the manuscript or revising it critically for important intellectual content. Preparing the final approval of the version to be published.

